# Adaptive evolution can overwrite initial natural fitness variation only in highest-fitness yeast isolates

**DOI:** 10.1101/2025.10.09.681448

**Authors:** Alexandra N Khristich, Olivia M Ghosh, Jean CC Vila, Shaili Mathur, Abhishek Dutta, Marion Garin, Joseph Schacherer, Dmitri A Petrov

**Author notes:** contributed equally (Alexandra N Khristich, Olivia M Ghosh).

## Abstract

The fitness of an organism determines its likelihood of succeeding in short-term competition, but many other factors can influence its long-term success. In this study, we investigate the relative contribution of initial fitness to a strain’s long-term success in a naturally diverse population. Specifically, we compete a pool of 282 genetically barcoded *S. cerevisiae* isolates for up to 720 generations in six distinct environments. We find that the strains that remain at detectable frequency until the end of the competition uniformly come from the initially fittest top 5% of strains, rendering initial fitness a strong predictor of long-term success. However, small fitness differences among the top strains matter little for their long-term evolutionary fate. Indeed, we often see heterogeneity in the competition outcomes across replicates, suggesting that stochastic, large-effect adaptive mutations overwrite small differences in the initial fitness of the highest fitness strains. In addition, our results hint that evolvability differences might result in certain strains consistently over- or underperforming in the competition. We further demonstrate that the “finalists” of our competition accumulate a diverse spectrum of *de novo* genetic changes: single nucleotide mutations, indels, whole chromosome losses and amplifications, and widespread losses of heterozygosity that occasionally span hundreds of kilobases. Taken together, we show that for a pool of natural strains, high initial fitness is necessary, though not sufficient, to succeed long-term, and that adaptive evolution can drive unexpected outcomes in a long-term competition in a novel environment.

## Introduction

A central goal of evolutionary biology is to predict the probability of a given genetic lineage’s long-term evolutionary success. Having a high current fitness is clearly important for short-term success and gives two distinct advantages in the long-term. First, by definition, a lineage with higher relative fitness expands in frequency and is more likely to outcompete other genotypes. But by increasing in frequency, a fitter lineage also increases its capacity to produce beneficial mutations due to its larger population size, and thus has a head start in the evolutionary competition (Desai and Fisher 2007; Fisher 2013). Indeed, microbial evolution experiments have demonstrated that in evolving clonal populations, lineages that happen to acquire adaptive mutations first outcompete others, generating a predictable statistical pattern colloquially referred to as ‘rich get richer’ (Levy et al. 2015; Blundell et al. 2019; Nguyen Ba et al. 2019).

However, in evolving populations, adaptive mutations arise fundamentally stochastically and can have very large effect sizes (Levy et al. 2015), so that even low- or medium-fitness genotypes can at times ‘leapfrog’ over higher-fitness counterparts (Kao and Sherlock 2008; Gerrish and Lenski 1998). This means that for a population where the magnitude of adaptive leaps exceeds the magnitude of initial fitness variation, initial fitness is a weak predictor of long-term success. In such populations long-term population composition will be hard to predict.

Moreover, some lineages might have a higher inherent ‘evolvability’, defined as an ability to acquire large adaptive mutations at a higher rate than others (Woods et al. 2011). Theory suggests that such high-evolvability lineages would tend to “win” in the long-term, even despite possibly lower initial fitness (Kirschner and Gerhart 1998; Earl and Deem 2004; Ferrare and Good 2024). In an extreme case, where variation in evolvability dramatically exceeds variation in starting fitness, the outcome of evolutionary competition will be fully determined by evolvability. While the magnitude of variation in evolvability is not known, previous studies have provided empirical evidence that this variation does exist in natural populations. Indeed, both the distributions of fitness effects (DFE) and mutation rate depend on genetic background, and vary significantly even within species (Kryazhimskiy et al. 2014; Gou et al. 2019; Jiang et al. 2021; Woods et al. 2011; Jerison et al. 2017).

Obviously, determinants of long-term outcomes in an evolution experiment will depend on the nature of the experimental system. Take a population of previously evolved adaptive mutants with dramatic fitness differences, and the long-term composition will likely be determined by the initial fitnesses alone (Nguyen Ba et al. 2019). Take a clonal population, and the long-term composition will be determined solely by stochastic adaptive mutations (Levy et al. 2015; Kao and Sherlock 2008). Take strains with engineered mutation rates, and the long-term composition could be determined by their evolvability differences (Gibson et al. 1970; Tröbner and Piechocki 1984, 1981; Chao and Cox 1983; Gentile et al. 2011). But what happens if we take a natural population? To answer this question, we ran a long-term competitive evolution experiment with a collection of natural *S. cerevisiae* isolates (Dutta et al. 2026). The collection is representative of the genetic and ecological range of the species, and likely spans much of the standing phenotypic variation. This makes it a great model system in which initial fitness differences, magnitude of adaptive mutations, as well as variation in any traits possibly associated with evolvability reflect the values relevant to an evolving species in nature.

More specifically, in this study we compete a pool of 282 genetically barcoded natural yeast isolates for roughly 700 generations. We do so in six different environments, with 20 biological replicates per environment. Note that we did not choose our environments to represent any of the environments that yeast occupy in nature. Instead, this experiment mimics species’ introduction to a novel environment, and numerous previous experiments indicate that yeast adapt quickly to various laboratory batch culture environments (Levy et al. 2015; Kinsler et al. 2024; Kao and Sherlock 2008; Abreu et al. 2024). In addition, the high number of biological replicates allows us to determine how often the fittest strain wins, and the extent of genetic parallelism in adaptation. Finally, the six distinct environments ensure that the same genotype starts with different fitness in different environments.

What can we expect to see as a result of our experiment? Perhaps the standing genetic variation in initial fitness is so substantial that it completely determines the outcome of our competition. In this scenario, short-term fitness would be a perfect predictor of long-term success, and the initially fittest strain would always take over the population. But perhaps adaptive mutations arise so rapidly that they completely overwrite the initial fitness variation (Levy et al. 2015). In this scenario, we would see that the effects of the initial fitness are quickly erased by a constant reshuffling of the strains’ rank order and stochastic leapfrogging. In this case, the final pools will show great variability replicate to replicate, and will contain a variety of strains, except for extremely low-fitness ones that get lost immediately. Finally, perhaps existing variation in traits related to evolvability, such as ploidy (Zhu et al. 2016; Peter et al. 2018; Sharp et al. 2018), mutation rate (Gou et al. 2019; Jiang et al. 2021), heterozygosity levels (Peter et al. 2018; Dutta et al. 2021; Marsit et al. 2021), as well as possible differences in the size of adaptive targets, completely overwhelms differences in initial fitness between strains. This will lead to certain genotypes with favorable combinations of “evolvability” traits predictably adapting faster and taking over the population, regardless of their initial fitness. In this case, we would expect to see that some strains would acquire adaptive mutations quickly and persist until the end of the experiment, regardless of their initial fitness.

We find that reality lies somewhat in between the spectrum of possibilities described above. Our strains exhibit substantial variation in fitness in any environment tested, and the initial fitness is, in broad strokes, an excellent predictor of success in this pooled setting. The strains that succeed in our experiment are always drawn from the top 5% of strains, graded by initial fitness in each environment. However, within this high-fitness tail of the distribution, we often find that the outcome of the evolutionary competition varies between the replicates of the same environment, and some of the high-fitness strains never persist through the end of the experiment. Our results show that stochastic adaptation plays a large role and produces heterogeneity in the competition outcomes, and that variation in traits associated with evolvability might matter for long-term success amongst strains that are already very fit. We also discover that natural strains’ repertoire of genomic adaptation is much more diverse than in haploid laboratory strains, and consists of widespread losses of heterozygosity (including conversion to a minor allele in a triploid strain), chromosomal abnormalities, structural variation, as well as indels and point mutations. We see examples of convergence at the gene level both across evolution environments and strain backgrounds. Taken together, our results show that while adaptation from standing genetic variation is predictable at a coarse-grained level, additional mutations can lead to idiosyncratic “adaptive leaps” that are hard to predict.

## Results

### An experimental system to study the relationship between short-term fitness and long-term evolutionary success in a genetically diverse population

Here, we set up a long-term competition experiment using a collection of genetically barcoded wild yeast isolates (Dutta et al. 2026). This collection consists of 282 strains and encompasses much of the phylogenetic, geographic, and ecological diversity of the well-characterized 1011 yeast isolates collection (Fig. S1) (Peter et al. 2018). We excluded haploid strains to prevent mating between different isolates, strains with killer virus activity to preclude toxin secretion (Chan et al. 2024), and flocculating strains to minimize cell clumping. In addition to the wild isolates, each replicate of our experiment contained 6 barcoded clones of the commonly used haploid S288C lab strain as a reference strain and to estimate measurement noise. To initiate the evolution experiments, we first recovered barcoded pools from the freezer by growing them in YPD for 24 hours and preconditioning them in the corresponding evolution media for 48 hours. We propagated barcoded strain pools in 10 mL batch culture with 1:250 dilution every 2 days in 12-well reservoirs without shaking for up to 91 transfers (∼720 generations) (Fig. 1A). We used six different environments (Fig. 1A, Table 1), selected to span environments with a wide range of initial fitness values (Fig. 1C). We had 20 biological replicates per environment to ensure that we sampled the highly probable outcomes of the competition.

**Fig. 1.**
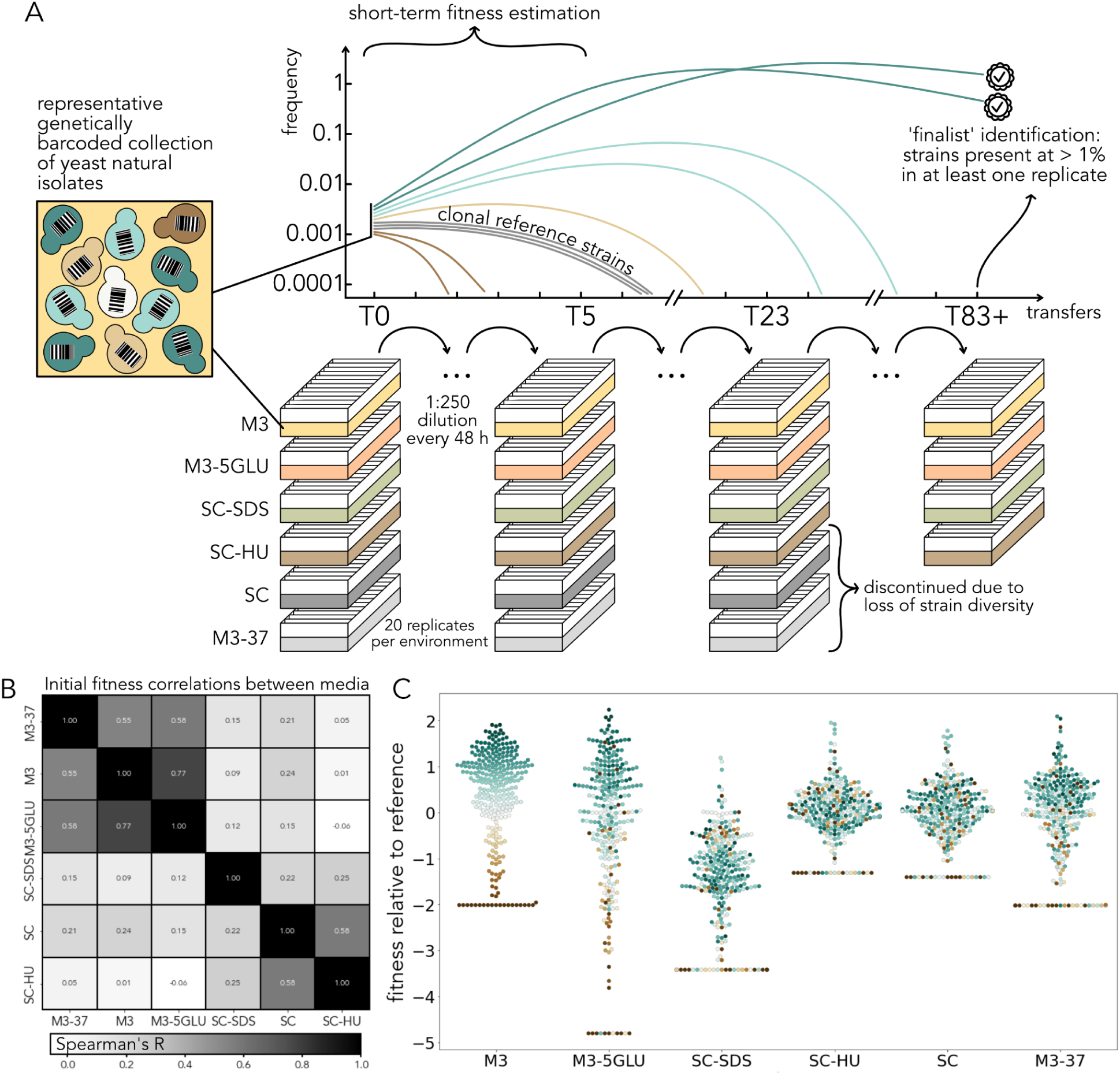
Experimental system to study the relationship between short-term fitness and long-term evolutionary success. **A.** A representative collection of 282 barcoded wild yeast isolates was propagated in a 10 mL batch culture for up to 91 48-hour transfers in 6 different environments, with 20 biological replicates per environment. Two of the environments were discontinued after 23 transfers due to rapid convergence to a single strain. Strain frequency trajectories from the first five transfers were used to compute short-term fitness, and frequency at the last transfer to identify the ‘finalists’ of the evolutionary competition. **B.** Spearman rank order correlation coefficient between fitness values of strains between pairs of environments. **C.** Distributions of relative fitness values in our environments. The colors represent relative fitness in M3.

**Table 1.** Environments used in this project.

| environment<br>nickname | base<br>media | glucose<br>% | incubation<br>temperature (°C) | added drugs | replicates | last sequenced<br>transfer |
| --- | --- | --- | --- | --- | --- | --- |
| M3 | M3 | 1.5 | 30 | none | 1-10 | 83 |
|  |  |  |  |  | 11-20 | 75 |
| M3-5GLU | M3 | 5 | 30 | none | 1-10 | 91 |
|  |  |  |  | none | 11-20 | 15 |
| SC-SDS | SC | 2 | 30 | sodium dodecyl sulfate<br>(SDS, 0.0075% final<br>concentration) | 1-10 | 88 |
|  |  |  |  |  | 11-20 | 79 |
| SC-HU | SC | 2 | 30 | hydroxyurea (HU, 3<br>mg/mL final<br>concentration) | 1-10 | 89 |
|  |  |  |  |  | 11-20 | 79 |
| M3-37 | M3 | 1.5 | 37 | none | 1-20 | 23 |
| SC | SC | 2 | 30 | none | 1-20 | 23 |

By amplicon sequencing the barcode locus, we constructed barcode frequency trajectories from the first 5 timepoints (Fig. 1A). We used these trajectories to infer the fitness of each of the natural strains relative to the lab reference strain using a previously developed maximum likelihood approach (Ascensao et al. 2023; Ghosh et al. 2026). Fitness values of the same strain varied dramatically between the environments, with some environments being practically uncorrelated, and some, for example, M3 and M3-5GLU, having similar distributions of initial fitness values (Fig. 1B,C). The shapes of the tails of the fitness value distributions also varied across environments, with some environments having many high fitness strains and others, such as SC-HU, having just four strains that do not immediately go extinct in this media.

In two environments, M3-37 and SC, the initially diverse population lost almost all strain diversity in all 20 replicates within the first 5 transfers, with just one strain dominating in each of the replicates (CGB and BAN, respectively). These strains were initially amongst the fittest in the population in their corresponding environment and started the competition at a high frequency. We confirmed that in each of these two environments, the same strain remained dominant up to 23 transfers, or 184 generations, at which point we discontinued these environments (Fig. S4, S5).

### Initial fitness is a strong predictor of long-term success

At the end of this 6-month experiment, we sequenced the last timepoint for each of the replicates of the remaining four environments, as well as several timepoints from the middle of the experiment. Due to unforeseen technical issues, the last timepoint that was available for sequencing varied among replicates within each environment (Table 1), and for the M3-5GLU environment, we only had 10 replicates that survived through the end of the experiment. The overall diversity in the pool composition declined at varying speeds in different environments, and the most dramatic changes occurred in the first 10 transfers (Fig. 2A, B, S2-7). Next, we recorded the strains that persisted at high frequency in each environment, and their resulting frequencies within-replicate and within-environment. We define a strain as a “finalist” in a given environment if it was present at ≥1% frequency in at least one replicate at the last available timepoint. No environment had more than five finalists. The SC-SDS environment had the same single finalist in 19 out of 20 replicates, and the M3, M3-5GLU, and SC-HU environments had between 3 and 5 finalists, which ended up in different proportions at the final timepoint (Fig. 2E). Altogether, the long-term dynamics in our experiment revealed a variety of qualitatively different outcomes: clear dominance of a single strain in SC-SDS, M3-37, and SC, possible long-term coexistence of the same three strains in M3-5GLU, alternative states with one or two dominant strains in SC-HU, and a mix of all the above outcomes in M3 (Fig. 2E).

**Fig. 2.**
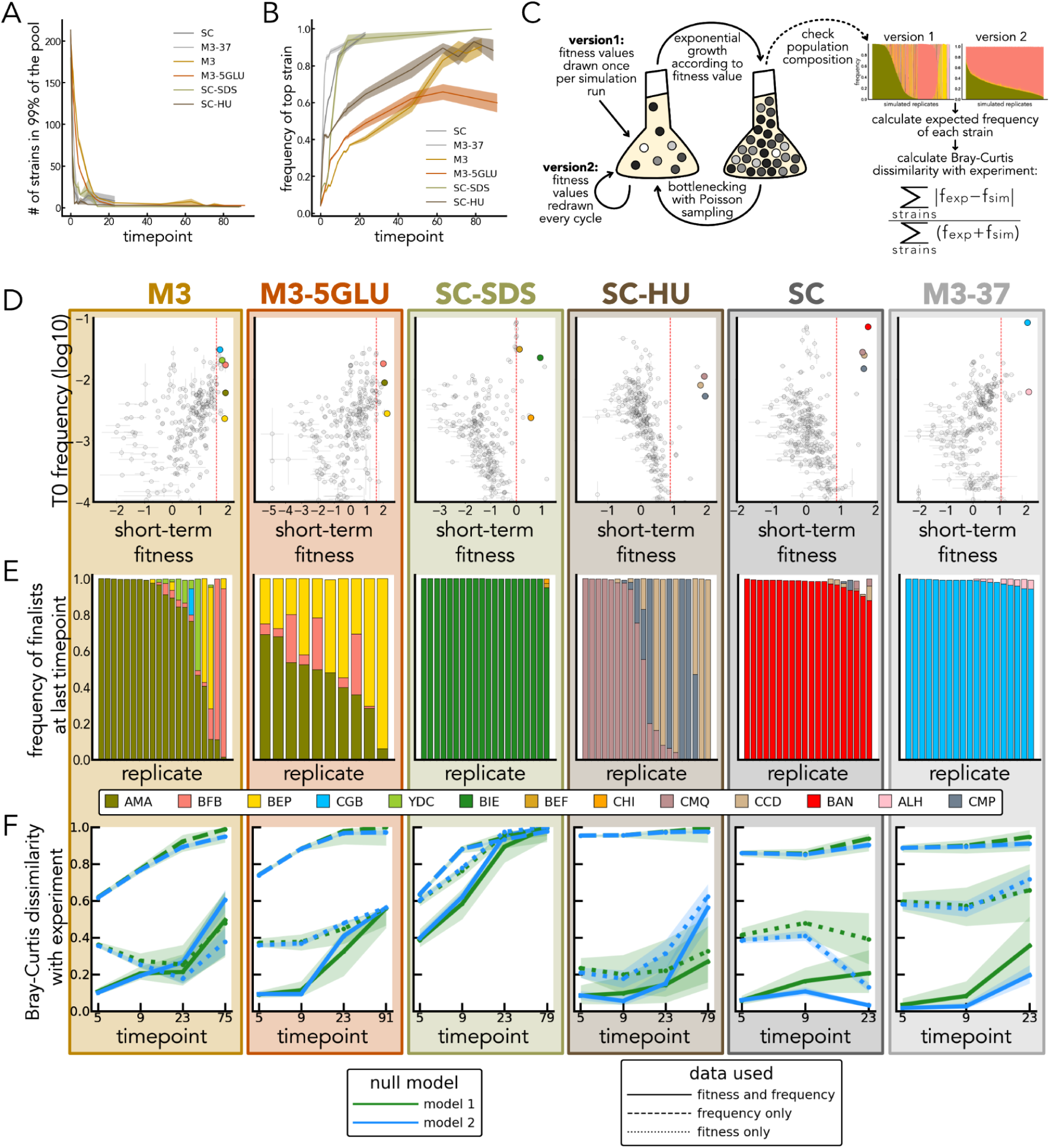
The finalists of the evolutionary competition. **A.** Overall dynamics of the pool composition in each environment. The median number of strains composing the majority of the pool (99%) at different timepoints of the long-term evolution experiment and their interquartile ranges. **B**. Frequency of the most abundant strain over time in each environment and their standard deviations. **C.** Simulating the two null models. In both null models, relative fitness values are drawn from a normal distribution centered at the estimated fitness value for each strain with the spread equal to the estimated error. In version 1, fitness values were drawn once per simulation run, and in version 2 once every cycle. The average frequency of each strain at selected timepoints is compared with experimental data. The simulated data for timepoint 75 from M3 media is shown as an example (to be compared with Fig. 2E, M3 panel). **D.** Relative short-term fitness calculated from the timepoints T0-T5 using the maximum likelihood framework and mean frequency at T0 for all strains in the pool. Colors other than light grey indicate finalist strains. The error bars represent the inferred standard errors. Red dashed lines represent 95th percentile in fitness values. **E.** Experimental frequencies of all strains present at ≥ 1% at the last available timepoint for each replicate. Colors are consistent with the colors from other panels. **F.** Comparing final average frequencies of the natural strains at the last timepoint of the evolution competition between the experiment and the null models.

In the simplest case, without evolution or frequency-dependent interactions, we reasoned that the long-term success of a strain is determined purely by its initial frequency and fitness. Note that while we aimed to start all of the strains at the same frequency, the experiment begins after propagation of the pools, a freeze-thaw cycle, and pre-conditioning in the evolution media, leading to unavoidable variation in initial strain frequencies. Therefore, we calculated the mean frequency of all strains at T0 and inferred their fitness from the first five transfers of the competition using the maximum likelihood method. While the initial frequency of the finalists varied, they were always in the top 5% of initial fitness in their respective environments (Fig. 2D). Interestingly, several strains in the top 5% of fitness values did not persist at high frequency to the end of the experiment in any replicate.

Next, we tested whether the observed heterogeneity in the competition outcomes and aforementioned underperforming strains could be explained by a simple exponential growth model with sampling noise (Fig. 2C). Because our strains exhibit substantial variation in initial fitness and frequencies, we tested whether a null model that takes into account strains’ starting fitnesses, starting frequencies, or both, can explain the competition outcomes. We simulated exponential growth of each strain in each of the media, using its starting frequency at T0 and estimated fitness in the corresponding media. Each of our fitness values comes with uncertainty, which is composed of the measurement error and biological noise between the 20 biological replicates. To account for this uncertainty, we drew each strain’s fitness from a normal distribution centered at its estimated fitness value, with the spread reflecting our inferred error. In the first null model, we drew fitness once per simulation run, and in the second null model, during every transfer (Fig. 2C). After 1000 iterations of the simulations, we bootstrapped simulated replicates (to match the number of experimental replicates for each media type) and compared the averaged composition of the final pool to the experimental data. In agreement with the experimental data, both null models revealed heterogeneity of outcomes at the last timepoint of the competition. However, the strain abundances in the simulated pools differed from those in the experiment (Fig. 2E, S8).

To quantitatively compare the population composition in the null models with the experiment, we first calculated the average frequency for each strain at selected timepoints of the competition. We then calculated the Bray-Curtis dissimilarity between these average frequencies for our experimental data and the bootstrapped simulation runs (Fig. 2F). We found that while the replicate-to-replicate strain composition differed between the two null models, the average frequencies of the strains were fairly similar between the two models (Fig. S9). The null models built on initial frequencies alone (assuming all strains are neutral with a measurement error) failed to explain the data. Null models built using both fitness and initial frequencies predicted strain composition well in the early timepoints (except for the SC-SDS environment), but the prediction strength was uniformly reduced by the last timepoint. The poor alignment between the data and simulation in SC-SDS might be explained by the fact that the fittest strains in this media violate exponential growth assumptions even in the first five transfers, as evidenced by their U-shaped frequency trajectories (Fig. S7). This could be due to long-lasting ‘memory’ effects arising as a result of transitioning from pre-growth media (Abreu et al. 2024), or possible ecological interactions between the strains at the beginning of the evolutionary competition. For the other environments, our results suggest that initial fitness is a strong predictor of long-term population composition. However, as time goes on, the predictive power of initial fitness decreases. In all cases, by the end of the experiment, a simple null model with a single fitness value per strain fails to quantitatively capture the resulting distribution of strain frequencies, suggesting a role for more complicated effects like time-varying selection or stochastic adaptation.

To explore the possibility of stochastic adaptation in more detail, we focused on the trajectories of the finalists in M3 (Fig. 3D, S2), as this media had the most diverse outcomes in the long-term competition. After 23 transfers (approximately 184 generations), only 7 strains are present in at least one replicate at a frequency above 1%, five of which ultimately become the finalists (AMA, BEP, BFB, CGB, and YDC). After about 30 transfers, the strain frequencies start diverging (Fig. 3B, S2) and occasionally flip in rank order, which ultimately leads to the final heterogeneity of outcomes.

**Fig. 3.**
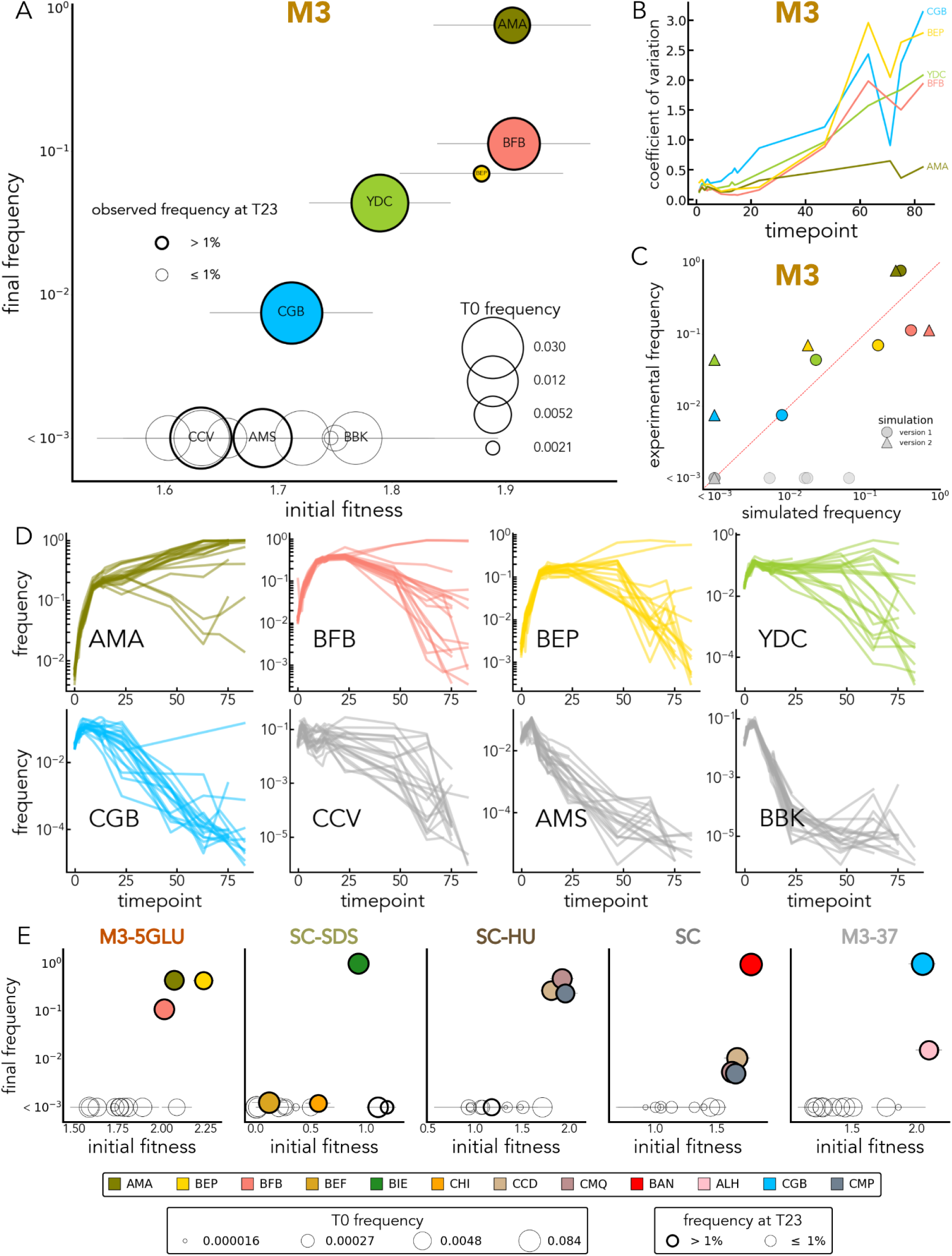
Among the fittest strains, initial fitness does not perfectly predict the end frequency. **A.** The initial fitness and final frequency at the last available timepoint for the top 5% fittest strains in M3. The finalist strains are colored by their strain identity, and the sizes of circles reflect their starting frequency. Other strains from the top 5% fittest strains (these strains never reach 1% frequency in any of the replicates at the last timepoint and their final average frequency is always < 0.003) are shown but not colored. Faint grey lines represent our error estimates. **B.** Coefficient of variation across experimental replicates in the M3 finalist strain frequencies over time. **C.** Final frequencies of the top 5% strains in M3 in data and simulated null models. **D.** Trajectories of the finalist strains in M3, as well as 3 other fittest strains in all replicates. **E.** The initial fitness and final frequency of the top 5% fittest strains in the other five media types. Sizes of the dots reflect their initial frequency for the corresponding media. Faint grey lines represent our error estimates.

### Frequency dynamics during long-term competition suggest a role for adaptive evolution

Interestingly, the initial relative fitness (calculated from the first five transfers) of the top 5% fittest strains appears to be a poor predictor of their relative abundance in the final pools. First, there are four strains initially more fit than the CGB strain that never reach detectable frequency past the first 40 transfers (Fig. 3A). For example, the BBK strain is the fifth fittest strain in M3, and it stays below 1% frequency starting from the 9th transfer, unlike CGB, which is the ninth fittest strain and one of the eventual finalists. Another interesting example is CCV, the 13th fittest strain, which remains at frequencies above 1% up to the 63rd transfer, but eventually decreases in frequency. Of the strains that do remain and become finalists, BFB has the highest fitness and initial abundance (Fig. 2D, 3A). Yet, compared to the simulated null models, it dramatically underperforms in the long-term competition, especially compared to the AMA strain, which dramatically overperforms (Fig. 3C). We observe similar results in most of the other environments, where the small differences in the initially high-fitness strains appear irrelevant as the evolutionary competition goes on (Fig. 3, S11). As a result, the final strain frequencies do not align with predictions of the null models in M3-5GLU, SC-HU, M3-37, and SC-SDS (Fig. 2F, 3E, S10), although the SC-SDS environment is slightly complicated by the fact that the exponential model is a poor fit for initial strain fitnesses (Fig. S7).

In addition, we occasionally observe what appears to be the effect of large adaptive mutations which result in a dramatic increase in frequency of specific strains (Fig. 3D, S12-14). In M3, for example, the most striking examples of such apparent adaptive mutations are evident for several replicates of BEP and BFB strains (between transfers 20 and 40) and for one replicate of the CGB strain (between transfers 30 and 50) (Fig. 3D, S2). In support of the idea that our yeast strains have evolved adaptively, pairwise competitions between the evolved clones and a fluorescently labeled strain confirmed that most of the finalists of our experiments outperform their unevolved ancestors (Fig. S19).

Taken together, a closer inspection of the frequency dynamics in our competition revealed hidden complexity behind a simple competition experiment (Fig. 3). Among the top fitness strains, stochastic forces, likely adaptation, appear to shape the outcomes of a long-term competition, rather than initial rank within the high-fitness class. This is consistent with a model where stochastic large-effect adaptive mutation and clonal interference determine the evolutionary fate of a complex population (Kao and Sherlock 2008).

### Whole genome sequencing of evolved isolates reveals a diverse repertoire of *de novo* genetic changes

To confirm that evolution happened at the timescale of our experiment, we used whole-genome sequencing to identify *de novo* mutations that occurred and rose to detectable frequencies during the course of our 6-month competition. Unlike haploid or homozygous diploid lab strains, typically used for experimental evolution research (Levy et al. 2015; Nguyen Ba et al. 2019), some of the finalist strains in our experiment were heterozygous diploid or triploid strains. Thus, it is important to understand what kind of molecular changes drive adaptation of natural strains, with this additional complexity in genome architecture. Therefore, we isolated and whole genome sequenced evolved clones at their last timepoint in each of the full-term environments (Table S5). We detected substantial private genetic variation in each evolved strain (Fig. 4A). The types of *de novo* mutations varied widely among the sequenced finalist isolates and included aneuploidies, large structural variants, novel heterozygous and homozygous point mutations, and small insertions and deletions (indels) (Fig. 4B,C, S15). In addition to *de novo* mutations, we also detected widespread loss of heterozygosity (LOH) events in the heterozygous strains (Fig. 5).

**Fig. 4.**
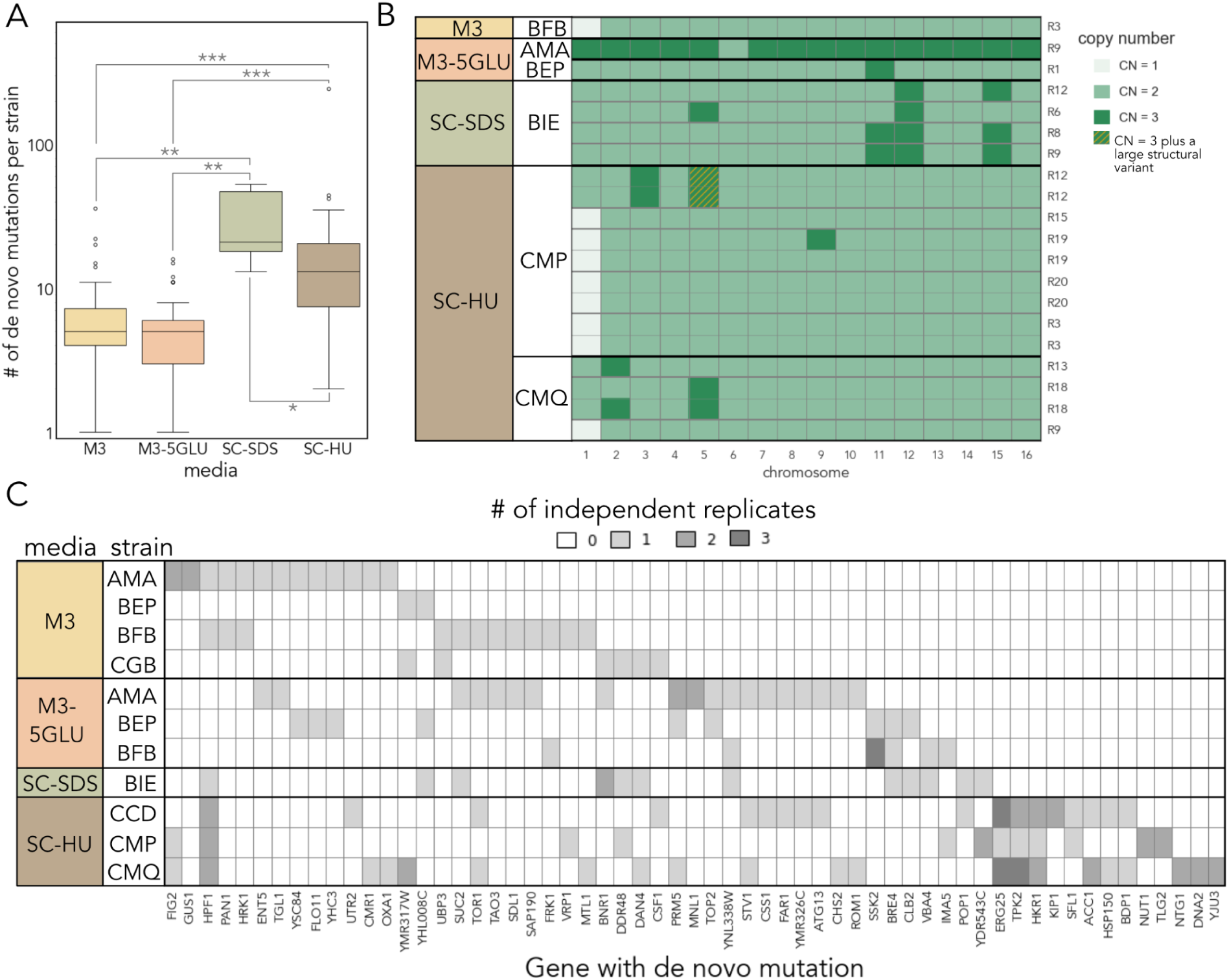
*De novo* genetic variation in the evolved clones. **A.** Boxplots showing *de novo* mutations per strain isolated from each evolution media. Asterisks refer to P-values obtained using the Mann-Whitney U test and adjusted for multiple comparisons using the Bonferroni correction. Specifically *: P-value < 0.05, **: P < 0.01, ***: P-value < 0.001. **B.** Summary of *de novo* aneuploidies and large structural variation detected in evolved clones. Only strains with aneuploidies detected are shown. Thick black horizontal lines separate strains evolved from the same ancestor and within the same environment; medium grey horizontal lines separate strains derived from independent replicates of the same evolution media; thin light grey lines separate individual strains derived from the same evolution replicates (thus might not represent independent events). Note that AMA strain is a triploid; thus the AMA clone on this plot has a loss of one copy of chromosome 6. **C.** Examples of parallel evolution in our experiment. The heatmap shows genes in which *de novo* SNPs or indels that alter protein-coding sequences arose more than once in our evolution experiment.

**Fig. 5.**
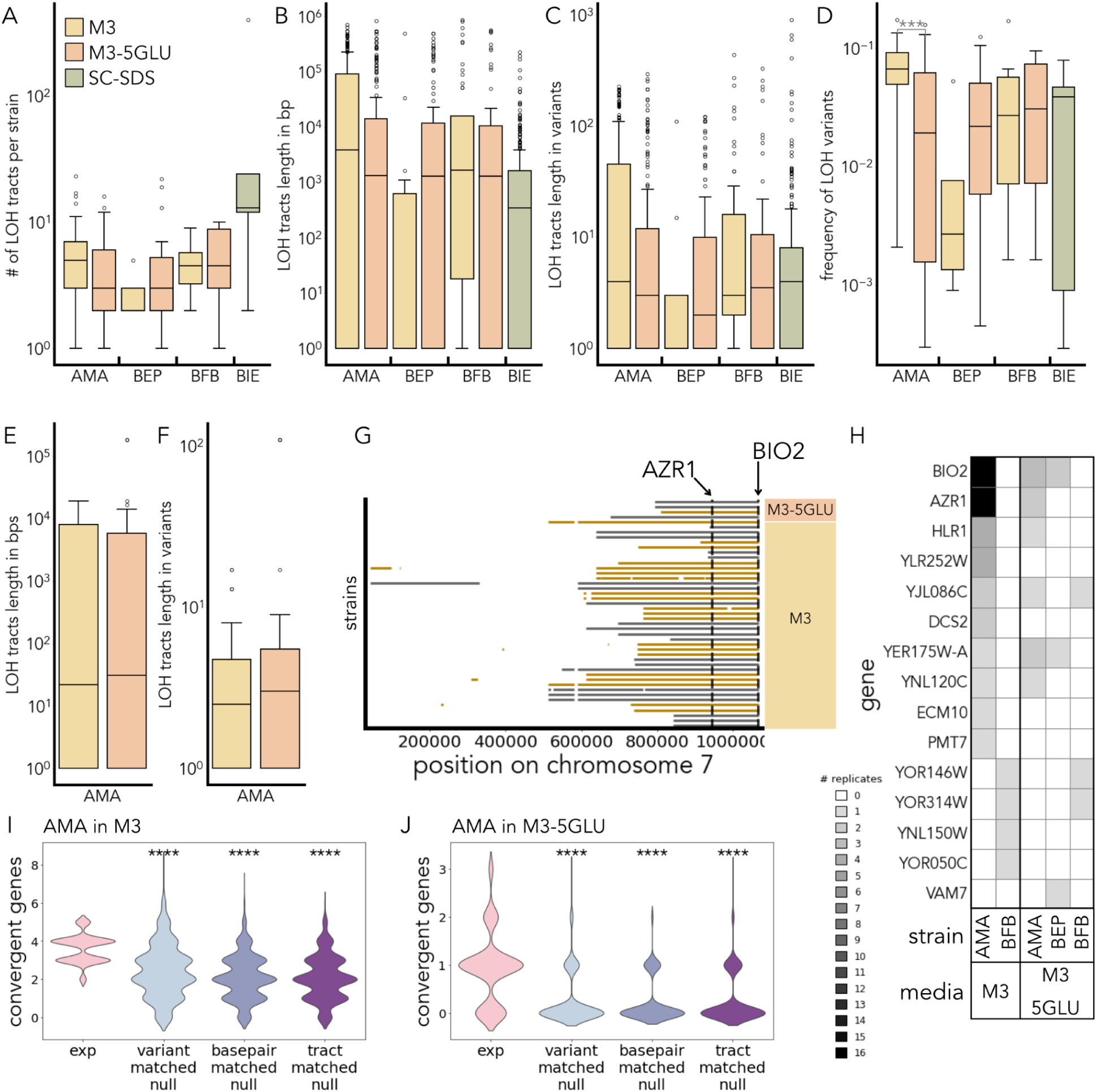
LOH events in the evolved clones. **A.** Boxplots showing distribution of the total number of independent LOH tracts per evolved strain as a function of strain background and evolution media. **B.** Boxplots showing distribution of lengths of independent LOH events, measured in basepairs, from all strains isolated from a given strain background and evolution media environment. **C.** Boxplots showing distribution of lengths of independent LOH events, measured in a number of converted variants (initially heterozygous variants that became homozygous), from all strains isolated from a given strain background and evolution media environment. **D.** Boxplots showing distribution of the frequency of heterozygous variants, converted to a homozygous state, per evolved strain as a function of strain background and evolution media. ***: P-value < 0.001 (Mann-Whitney U test) **E.** Boxplots showing distribution of lengths of independent LOH events, consisting of heterozygous variants converted to the minor allele, measured in basepairs, from all triploid AMA-derived strains isolated as a function of evolution media. **F.** Boxplots showing distribution of lengths of independent LOH events, consisting of heterozygous variants converted to the minor allele, measured in a number of variants, from all triploid AMA-derived strains isolated as a function of evolution media. **G.** LOH tracts from chromosome 7 from the strains with a conversion to the frameshift variant in the AZR1 gene. Each row on the y axis represents an individual finalist strain. The strains are ordered first by evolution media, and then by evolution replicate. Colors of the lines are used to separate strains from independent replicates (two adjacent strains in the same color might not be independent). **H.** Examples of parallel evolution in our experiment. The heatmap shows genes in which high-impact LOH variants converted to the same homozygous state more than once in our evolution experiment. **I, J.** Quantification of the number of convergent genes between AMA-derived strains isolated from independent replicates in M3 (*I*) and M3-5GLU (**J**) evolution media, as described in the text. Asterisks refer to P-values obtained using a one-sided Mann-Whitney U test, where an alternative hypothesis is that the central tendency of the null distribution is smaller than in the experimental one. ****: P-value < 0.0001.

The mutational spectrum depended on both the strain background and environment. For example, aneuploidies and large structural variants predominantly occurred in the SC-SDS and SC-HU environments (Fig. 4B). The aneuploidy events included both chromosomal losses and duplications, and 9 out of 16 chromosomes were observed in at least one aneuploidy event. The most common aneuploidy event was a loss of chromosome 1 that occurred in at least six independent evolutions across two evolution environments (SC-HU and M3) and three strain backgrounds. Most of the observed aneuploidy events happened in the SC-HU environment. This was likely driven by the mutagenicity of this environment: HU causes replication and oxidative stress in yeast (Shaw et al. 2024) and can trigger a variety of mutations, including point mutations, aneuploidies, and structural variants (J. Li et al. 2023). Surprisingly, SC-SDS also gave rise to a substantial number of aneuploidy events in the BIE strain, including recurrent amplifications of chromosomes 11, 12, and 15. Since BIE was the only finalist in SC-SDS that we isolated, it is impossible to know whether the high number of aneuploidies in it is due to the previously unknown property of SDS to trigger aneuploidy, high aneuploidy formation rate in the BIE strain, or the fitness benefit of specific aneuploidies in this environment (Sunshine et al. 2015). The first is unlikely because SDS is not a known mutagen. At concentrations 4 times higher than used here, SDS causes cell wall and oxidative stress (Cao et al. 2020; Zhao et al. 2020), but its ability to directly trigger chromosomal abnormalities has not been previously reported. The second is also unlikely as the rate of aneuploidies in the unperturbed BIE is low compared to other natural isolates (Dutta et al. 2022). Thus, the most likely cause for high prevalence of aneuploids among the BIE-derived strains is related to these aneuploidies’ positive fitness effects.

The total number of point mutations and indels per strain also depended on the evolution environment, with the SC-HU and SC-SDS media generating more mutations per strain than M3 or M3-5GLU environments (Fig. 4A, S15). Within each environment, we also observed a difference in the number of mutations per strain. The most pronounced differences were observed for the finalists of the SC-HU environment, where we detected statistically significant differences between the total number of all heterozygous mutations (Fig. S15A) and in total number of all point mutations (Fig. S15B) that arose in the three strain backgrounds.

We saw a few instances of parallel adaptation at the gene level across environments and strain backgrounds. Fig. 4C summarizes 59 genes where a *de novo* single nucleotide variant or indel of high or moderate impact arose in more than one independent evolution replicate. Every finalist strain background in every evolution environment has at least two genes with mutations that arose in more than one replicate. In addition, we saw examples of parallel evolution for different strains evolving in the same media: five genes in M3, five genes in M3-5GLU, and ten genes in SC-HU independently mutated in different finalists of each of these media. At the same time, we observe that the same two genes in AMA, one gene in BEP, and one gene in BFB mutated in both M3 and M3-5GLU environments. Taken together, our data suggest that both the evolution environment and strain background determine the evolutionary path.

### LOH events are the dominant mode of molecular evolution in heterozygous strains

The most common mutational event we observed in the heterozygous finalist strains (AMA, BEP, and BFB in M3 and M3-5GLU, and BIE in SC-SDS, Table S4) was LOH at ancestral heterozygous variants. In some of the evolved strains, up to 17% of the initially heterozygous variants converted to a homozygous state (Fig. 5D). The frequency of converted variants depended on the evolution environment for AMA, but not other strains (Fig. 5D). The converted variants are often clustered together alongside genomic coordinates in uninterrupted stretches we refer to as LOH tracts. Overall, in our experiment we observed an average of 8 independent LOH tracts per evolved strain (Fig. 5A). An average tract’s length was 43 kb in raw genome coordinates (Fig. 5B) or 23 variants (Fig. 5C). Many of the long LOH tracts terminated at the chromosomal end, which suggests that they might have been formed via the break-induced replication mechanism (Fig. 5G) (Wu and Malkova 2021). Strikingly, in the triploid AMA strain, a subset of LOH tracts formed via conversion to the minor allele (present in one copy at the initially heterozygous site) (Fig. 5E, F). Further mechanistic studies are needed to elucidate whether conversion to a minor allele in a triploid strain is a result of two independent mutational events, or if it could be produced in a single break-induced replication cascade.

Given the high prevalence and long tract lengths of LOH events in our evolution experiment, determining which of the individual LOH variants are functionally significant is not trivial. Therefore, we focused our attention only on protein-coding high-impact mutations such as frameshifts and nonsense mutations. We found several instances of the same loci converting to the same homozygous state in independent replicates of the same media, strain background, or both. The most striking examples include conversion to a loss of the start codon variant of the *BIO2* gene, and to a frameshift variant in the *AZR1* gene that happened 21 and 18 independent times, respectively, across two media types and two strain backgrounds (Fig. 5H). Both of these genes are located on the right arm of chromosome 7, and all LOH tracts with the *AZR1* variant also contained the *BIO2* variant (Fig. 5G). The *AZR1* gene is implicated in resistance to fluconazole and ketoconazole (Tenreiro et al. 2000), and a previous large-scale screen revealed that the absence of this gene provides a fitness advantage to yeast growing in minimal media (Breslow et al. 2008). Thus, it is possible that conversion to the frameshift *AZR1* variant is one of the beneficial mutations in the AMA background. On the other hand, losing the *BIO2* gene is likely neutral or nearly neutral. This gene is important for biotin metabolism, and biotin is already supplemented in the M3 and M3-5GLU media. It is also possible that the local chromosome architecture is responsible for the high incidence of LOH events in the region.

Since it is hard to identify driver mutations even among convergent LOH events, we decided to test whether we observe more mutations that converge at the gene level than expected by chance. For that, we first bootstrapped our data such that, in each bootstrap, we randomly sampled one strain from each replicate of evolution media and strain background and calculated the number of high-impact variants that independently convert to the same state across experimental replicates. Since we have the overlapping three finalists in M3 and M3-5GLU media, we quantified convergence on three levels: 1) within the same evolution media and strain background, 2) across two or more strain backgrounds within the same media, and 3) across two or more evolution media within the same strain background. We generated a null expectation for convergence at each of these levels and compared it to our experimental distribution. For each null model, we randomly converted a number of heterozygous variants per each strain such that the total number of converted variants matched the data for this strain (variant-matched null), the number of LOH tracts and their length in raw base pairs matched the data for this strain (base pair-matched null), or the number of LOH tracts and their length in variants matched the data for this strain (tract-matched null). We found that the experimental number of convergent genes in the replicates of the AMA strain evolving in M3 (Fig. 5I) and M3-5GLU (Fig. 5J), and for the BEP strain evolving in M3-5GLU (Fig. S16), significantly exceeds the null distribution irrespective of the null model used. We also found that convergence of LOH variants between different strains in M3-5GLU (Fig. S17) and between the two media types in the AMA strain background (Fig. S18) also exceeded any null expectation. Overall, these results strongly suggest that a fraction of LOH events are selected for.

Taken together, we found that the timescale of our competition allowed enough time for novel mutations to arise, and the number and spectrum of these mutations depended both on the environment and strain background. On the genetic level, we detected a few examples of convergence both for *de novo* aneuploidies, SNPs, and indels, as well as LOH variants. However, most of the repeated gene targets of adaptation appear to be specific to the strain-environment combination, which suggests that adaptive evolution picks up on small differences between both the genetic background and the environment that are invisible to selection on standing genetic variation.

## Discussion

In this study, we used a genetically barcoded pooled collection of wild *S. cerevisiae* isolates to track strains as they competed and evolved in real time, to test the determinants of long-term success in a novel environment for within-species competition. This collection of natural isolates likely encompasses differences in most variable traits within the species, including substantial variation in initial fitness in the six novel environments used here. This variation should also include any traits associated with evolvability. When a clonal yeast population adapts to a lab condition, adaptation typically happens rapidly and involves large-effect adaptive leaps (Levy et al. 2015; Kao and Sherlock 2008). By focusing on natural strains rather than clonal yeast, we asked if standing variation in initial fitness or adaptation via *de novo* mutations contributes more to the population dynamics in a pooled evolution competition.

We found that in all six of the evolution environments in our experiment, the standing variation in fitness per cycle ranged over an order of magnitude (Fig. 1C). Not surprisingly, a strain’s initial fitness was a strong predictor of its long-term success, as only strains from the 5% fittest tail in each of the environments persisted in the long-term (Fig. 2D, E). This means that when a population enters a new environment, selection immediately sorts genotypes based on the standing variation in fitness. In some of our environments, such sorting alone determined the outcome of the competition. For example, in SC, a single highest-fitness strain dominated the population in every experimental replicate by the 23rd transfer (Fig. 2, S5), and other strains were not able to ‘leapfrog’ over this strain.

However, several of our environments had multiple strains in the high fitness tail of the fitness distribution, all of which started the experiment at relatively high frequency. In such cases, we observed heterogeneity in the long-term competition outcomes (Fig. 2E, 3A,E). The strain frequency dynamics in these environments were consistent with the presence of evolutionary adaptation via *de novo* mutations. Among the high fitness strains, we observed rank order switching and high variability in the final pool composition. This suggests that during the evolutionary competition, high fitness strains acquire large-effect adaptive mutations, which render the initially small differences in their fitness irrelevant to long-term success. We found that a simple model built only on the strains’ initial fitness and frequencies accurately predicted the population composition at the early timepoints, but not at the latest timepoints, which suggests either time-dependent fitness or adaptation. Furthermore, whole-genome sequencing of the evolved clones revealed aneuploidies, large structural variants, point mutations, indels, and massive LOH tracts. We detected convergence for the gene targets of point mutations, indels, and LOH events, suggesting a role for positive selection.

While the frequency trajectories, whole-genome-sequencing, and fitness remeasurements were consistent with adaptive evolution being the driver of the heterogeneity and rank changes in the strains’ frequencies at the later stages of our experiment, we cannot formally rule out the contributions of subtle changes in media composition, batch variation, or emerging ecological interactions to the observed results. For example, in M3, BBK is the fifth fittest strain, but its frequency drops rapidly after the first 9 transfers. It is possible that this is because BBK’s fitness decreases with the changing composition of the pool. In one of our environments, SC-SDS, frequency trajectories of some strains are U-shaped, even in the first five transfers we used to estimate initial fitness. This might be due to ecological interaction between the competing strains, a possibility worth investigating further in future studies.

What do our results mean for the notion of evolvability? The collection of natural yeast isolates encompasses the natural diversity of this species, and exhibits documented variation not only in the fitness of the strains, but also in evolvability-related traits such as mutation rate (Gou et al. 2019; Jiang et al. 2021). They also likely differ in their adaptive DFEs (Kryazhimskiy et al. 2014), although this has not been measured directly for this collection. Our observations suggest that differences in evolvability might be important in some cases for determining the outcome of competition between the high-fitness strains. For example, some of the 5% fittest strains never become finalists in our competition, even though their initial position (high fitness and high frequency) is similar to those strains that do persist till the end. Another intriguing example is the CCV strain that persists at high frequency for up to 63 transfers, but never seems to acquire a large-effect adaptive mutation.

The diverse spectrum of *de novo* mutations that we detected in the evolved strains hints at the possible molecular basis for evolvability differences between the strains. Many of the finalist strains are heterozygous, and LOH events were the most common mutational event in the heterozygous strains. LOH events present a robust path for adaptation, as they are common, unmask existing variation already present in the genome, and can affect many loci in a single mutational event (Gerstein et al. 2014; C. S. Smukowski Heil et al. 2017; Lancaster et al. 2019; C. Smukowski Heil 2023). Consistent with this idea, the AMA strain, which overperformed the null models in M3 (Fig. 3C) and M3-5GLU (Fig. S10), underwent numerous and extensive LOH events in the course of the evolutionary competition. However, a key difference between AMA and other heterozygous strains is that AMA is a triploid, whereas all other finalists are diploid. It is tempting to speculate that LOH events in a triploid, heterozygous strain provide an opportunity for even more adaptive mutations than in a diploid strain. Having a ploidy of three instead of two might present more initial haplotype combinations that the genome can convert to, as well as allow fine-tuning of allele dosage.

This study opens up many interesting avenues for future research. For environments with high heterogeneity in the competition outcome, repeating the experiment with a smaller pool of initial strains and more replicates would generate high-resolution measurements of long-term success. By varying the starting frequencies of the strains, or by allowing them to evolve in isolation before pooling, we could test how frequency-dependent interactions (Ascensao et al. 2025) and priority effects (Stroud et al. 2024) shape the outcomes of a long-term competition. By performing the experiment in fluctuating environments, we could establish the role of *de novo* mutations in a situation where fitness values are constantly reshuffled (Abreu et al. 2024) due to environmental change. Finally, by carefully measuring adaptive DFE of selected strains, we could decipher whether the under- and overperformance of certain strains in our experiment is due to subtle differences in evolvability between strains or simply due to a stochastic adaptive process.

To conclude, barcoded natural yeast strains proved to be an excellent model system to study the role of standing genetic variation in evolutionary dynamics. Using this system, we demonstrated that initial fitness is a strong predictor of the chances of long-term success for a genotype, and that *de novo* mutations can reshuffle the initial fitness values only for the strains from the top part of the fitness distribution. Our results hint that for these strains, evolvability differences might be important for long-term evolutionary success. In addition, we showed that natural strains exhibit an incredibly diverse repertoire of genetic changes as they evolve, which makes them a great system to study molecular evolution in heterozygous genomes of various ploidy levels.

## Acknowledgements

We thank Dr. Michelle Hays for generously providing strains used in the killer assay and for guidance in adapting the assay protocol. We also thank Dr. Benjamin Good, James Ferrare, and members of the Petrov lab for helpful feedback and discussions. We thank the Stanford Research Computing Center for the use of computation resources on the Sherlock cluster. A.N.K. acknowledges support from NIH NIGMS Grant No. F32GM149046-02. O.M.G. acknowledges support from the NSF-GRFP. D.A.P. acknowledges support from NIH NIGMS Grant No. 5R35GM118165-07. D.A.P. is a Chan Zuckerberg Biohub - San Francisco Investigator.

## Materials and Methods

### Barcoding wild yeast collection

#### Collection overview

The wild yeast isolates used in this experiment (Table S3) are a subset of the 1011 wild yeast isolates collection (Peter et al. 2018). This subset is composed of strains with unique DNA barcodes integrated into their genomes, to enable pooled assays (Dutta et al. 2026). For our pooled evolution competition, we first excluded from the barcoded collection strains with duplicate genetic barcodes and strains that did not grow after thawing a glycerol stock. Additionally, we excluded haploid, flocculating, and killer strains from the barcoded wild isolates collection.

#### Removing haploid strains

Haploid strains are a minority in the wild isolates collection, and to prevent mating between different isolates we decided not to include any haploid strains in our experiment. To test for haploids, we used a modified flow cytometry and SYBR green protocol to measure DNA content (Dunham et al. 2015). Individual strains in the collection, as well as known haploid and diploid control strains grew in YPD to saturation overnight and then diluted and grew for ∼5 hours to exponential phase in deep-well 2mL assay blocks. At this point, cultures were pelleted and resuspended and incubated at room temperature for 1 hour in 1mL of 70% ethanol. Cells were then washed and incubated in 0.2 mg/mL RNAse A solution at 37℃ for 2-4 hours to digest RNA and then washed and incubated at 50℃ in 2mg/mL proteinase K for 1 hour. After digestion steps, cells were resuspended into FACS buffer (200mM Tris-Cl pH 7.5, 200mM NaCl, 78mM MgCl_2_). 200 *μ*l of SYBR green diluted 5000X was added to each tube and mixed vigorously to reduce clumping and stored in the dark where possible. Samples were then run on the flow cytometer. We recorded at least 10000 events per strain and used the ‘BL1-H’ channel to estimate DNA content in each of the cells. We normalized the values to known haploid and diploid strains and assigned each of the strain ploidy of 1, 2, or higher. We excluded all the strains that scored as haploids from our pooled collection.

#### Removing flocculating strains

To measure flocculation in the wild isolates, we used a modified protocol from (Yone et al. 2022). We first inoculated glycerol stock of each strain into 1mL of YPD media and grew them overnight in 96-well deep well plates at 30℃ with no shaking. Next day, we used the Integra Viaflo 96 to first mix the overnight cultures and transfer 500 *μ*l of the cells into a new 96-well deep well plate. Then, we mixed these cultures by pipetting up and down 30 times at the highest speed, removed the pipette tips and waited for exactly 1 min. Next, we transferred the top 100 *μ*l of the cells into a new 96-well plate. In 10 sec, we added 100 *μ*l of 10 mM EDTA to the bottom fraction, mixed 30 times, and transferred 100 *μ*l of the bottom-cell-fraction-EDTA mix into a new 96-well plate. Immediately after the transferring of the top and bottom fraction cells into their respective 96-well plate, we measured the optical density of these samples at 600 nm using a BioTek Epoch2 platereader. We then subtracted the blank media OD measurement from every sample and calculated flocculation score for each strain using the formula: flocculation score =1 - OD_top cell fraction_ / OD_bottom cell fraction_ ⋅ 0.8. We confirmed high flocculation status for some of the known flocculators in the collection and additionally discovered 4 more flocculating strains. We did not include any strain that was either labeled as a flocculant in the collection metadata or scored as a flocculant in our assay into our experiment.

#### Removing killer strains

To identify wild strains producing killer toxin, we screened all barcoded natural isolates for K1 killer toxin production using a lawn-based assay adapted from Crabtree et al. 2019. Assays were performed on YPD agar buffered to pH 4.6 with sodium citrate (final concentration 0.1M). After autoclaving, 10 mL of methylene blue solution (final concentration 0.003% w/v) was added per liter as a marker of oxidative stress to enhance halo detection. Agar (50 mL) was poured into single-well rectangular dishes to ensure compatibility with a 96-well pipetting robot. Barcode and Lawn strains were revived from –80 °C glycerol stocks by streaking onto Sabouraud agar plates. A single colony of each barcoded strain was inoculated into 1 mL of Sabouraud broth in 96-well plates and incubated for 48 h at 25 °C with shaking (200 rpm) to saturation. A known “super-killer” strain (GSY1057) was included in one plate as a positive control. Cultures were maintained at 25 °C because killer toxins are heat-labile. Lawn cultures were prepared by inoculating 5 mL of Sabouraud broth in 25 mL flasks, incubating for 48 h at 25°C, and diluting 1:50 before plating. For each assay plate, 500 µL of diluted lawn culture was spread evenly across the agar surface. The primary lawn strain was a killer-sensitive BY-background strain (S) from (Pieczynska et al. 2016). As controls, we included the K1 killer strain (K) as a resistant lawn and an uninoculated mock lawn to monitor contamination and assess isolate growth in the absence of competition. For the assay, all test cultures were diluted 1:100, and 5 µL of each strain was spotted onto plates containing one of the three lawn types. Spots were applied in alternating positions, leaving empty wells between samples to allow sufficient space for halo detection. Plates were incubated at 25 °C for 5 days, after which halo formation was scored by visual inspection. All strains grew on the mock lawn. Strains that produced visible halos on sensitive lawns were recorded as killer-positive.

#### Pooling the strains

To pool the selected strains, we first streaked each of the barcoded wild strains to single colonies on a YPD plate, inoculated one colony into liquid 1 mL YPD media in a 96-deep-well plate, sealed with a ‘Breath-EASIER’ seal (Cat#: BERM-2000), and grew till saturation for 2 days at 200 rpm shaking at 30℃. After that, we pooled equal volumes of each strain together, mixed, and saved at −80℃ as 500 *μ*l glycerol stocks.

#### Barcoding reference strain

To barcode the lab strain and make it compatible with the barcoded natural isolates collection, we first amplified a fragment from the plasmid carrying a HygR gene, pSK26 (plasmid map included in the supplement), using primers SK168 and SK169 (Table S1). The resulting fragment contained the *HYG* resistance marker flanked by 20 bp random barcodes and regions of homology to the *HO* gene. We then transformed haploid strain GSY145 (S288C mat alpha, *ho*) (Kao et al. 2010) with this fragment. To validate the transformation, we amplified the DNA from the hygR clones with primers SK144/145 (Table S1) and Sanger sequenced the resulting 291 bp fragment to identify the barcodes of the newly generated barcoded strains (Table S2).

#### Media

To make media for the long-term competition, we first prepared the base media, M3 or SC in batches of 12 L following the recipes below. Then, we added the required amounts of the add-on drugs as needed for the specific media type (Table 2). We filter-sterilized the solutions and pre-poured them into the 12-well plates for storage and contamination control. We stored all media at room temperature except for the SC-HU media, which we stored at 4℃. During the growth, we sealed the plates with a ‘Breath-EASIER’ seal (Cat#: BERM-2000) and incubated them at 30℃ without shaking.

**Table 2.**
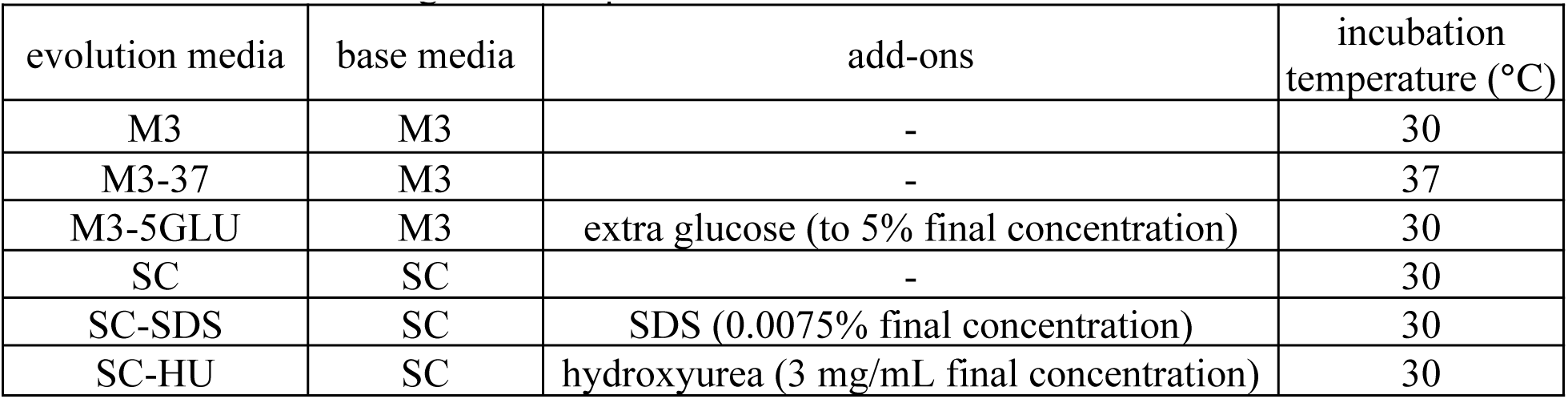
Media for the long-term competition.

| evolution media | base media | add-ons | incubation temperature (°C) |
| --- | --- | --- | --- |
| M3 | M3 | - | 30 |
| M3-37 | M3 | - | 37 |
| M3-5GLU | M3 | extra glucose (to 5% final concentration) | 30 |
| SC | SC | - | 30 |
| SC-SDS | SC | SDS (0.0075% final concentration) | 30 |
| SC-HU | SC | hydroxyurea (3 mg/mL final concentration) | 30 |

M3 (1L) modified from Verduyn et al. 1992:

- glucose: 15 g
- potassium phosphate monobasic: 15 g
- MgSO_4_*(H_2_O)_5_: 2.5 g
- ammonium sulphate: 40 g
- vitamin mix (1L: biotin 0.05 g, calcium pantothenate 1 g, nicotinic acid 1 g, inositol 25 g, thiamin HCl 1 g, pyridoxine HCl 1 g, para-aminobenzoic acid 0.2 g, 1000 mL H_2_O pH 6.5): 5 mL
- minerals mix (1L: EDTA (15 g EDTA, 4.5 g ZnSO_4_.7H_2_O, 1 g MnCl_2_*4H_2_O), 0.3 g CoCl_2_*6H_2_O, 0.3 g CuSO_4_*5H_2_O, 0.4 g Na_2_MoO_4_*2H_2_O, 3.4 g CaCl_2_, 3 g FeSO_4_*7H_2_O, 1 g H_3_BO_3_, 0.1 g KI, 1000 mL H_2_O, pH 4.0): 5 mL

SC (1L):

- YNB 6.7 g
- complete dropout 2 g
- glucose 20 g
- H_2_O 1000 mL

#### Evolutionary competitions

We first inoculated 500 *μ*l of glycerol stock containing the barcoded wild yeast pool into 80 mL of YPD media supplemented with 100 µg/mL carbenicillin to prevent bacterial contamination, and the barcoded reference strains into 5mL each and incubated for 1 day at 200 rpm and 30℃. Next day, we created subpools of reference strains (combinatorial combinations of strains which were unique for each replicate and media in our experiment) by mixing together equal amounts of each strain. Then, we added the wild yeast pool to each of the reference subpools in proportion 9:1 (wild yeast):(reference yeast). Finally, we transferred 40 *μ*l of the resulting pools with both wild and reference strains into 10 mL of the corresponding evolution condition media to initiate the timepoint 0 of our evolution experiment. We also saved aliquots of each of the pools in glycerol and sorbitol solutions for further DNA extractions.

From then on, we performed a transfer once every 48 hours by first carefully mixing the evolution media until the settled yeast became distributed evenly across the volume of the media. We then transferred 40 *μ*l of the cell suspension into fresh media. We saved a 100 *μ*l aliquot of the suspension in 16% glycerol solution at −80℃ and a 1mL aliquot in sorbitol solution (0.9 M sorbitol, 0.1 M Tris-HCL pH 7.5, 0.1 M EDTA pH 8.0) at −20℃ every 5 transfers, except for the first 38 transfers when we saved an aliquot in sorbitol every transfer. We continued the experiment for between 23 and 91 transfers, depending on the evolution media.

#### DNA extractions

We extracted DNA from 96-well plates using a custom column purification method. Briefly, we first thawed sorbitol-saved cells, spun them down, and discarded the supernatant. Next, we resuspended the cell pellets in 400 *μ*l tissue lysis buffer (NEB, cat# T3011L) supplemented with 7*μ*l of 10 mg/mL RNase A (Sigma, cat# R5000-1G/50-177-9327) and 1.2 *μ*l of 20 U/*μ*l lyticase (Sigma, cat# L4025-1MU), transferred the lysis buffer-cell mix to a deep well plate prefilled with 100 *μ*l of 500 *μ*m acid-washed glass beads (cat# G8772-1KG), and incubated at 37℃ overnight. The next day, we vortexed the lysis buffer-cell-bead mix for 5 min at the highest speed in a bead-beater, incubated at 65℃ for 15 minutes, spun down the plates for 10 min at 3700 rpm, and transferred the supernatant to a DNA purification column (cat# BD96-01). We washed the column twice with a wash buffer (NEB, cat# T3015-2) and eluted the extracted DNA into 40 *μ*l of elution buffer (NEB, cat# T3016-2).

#### Amplicon library preparations

To amplify the barcode for next-generation sequencing, we performed a two-step PCR protocol. For the first PCR step, we mixed 1 *μ*l of extracted DNA, 0.75 *μ*l of the forward and reverse 1st step primers at 10 *μ*M concentration (Table S1), 7.5 *μ*l of the NEBNext® Ultra™ II Q5® Master Mix (cat #M0544X), and 5 *μ*l of ultrapure water (Invitrogen, cat # 10977-015). The first step primers include a variable index region which allows for demultiplexing of the samples. We run the following PCR program:

Lid: 105℃, volume: 15 *μ*l

1. 98℃, 30 sec
2. 98℃, 30 sec
3. 66℃, 30 sec
4. 72℃, 30 sec
5. go to step 2 and repeat 35x times
6. 72℃, 5 min
7. 4℃, hold

After validating the correct size of the samples and the absence of amplification in the no-template control, we performed the second PCR step. First, we diluted the 1st step PCR product by 1000 fold using the ultrapure water. We then mixed 6 *μ*l of this diluted PCR product, 0.75 *μ*l of the forward and reverse 2nd step Illumina primers (Dual Indexed i5 and i7 primers), and 7.5 *μ*l of the master mix (NEB, cat #M0544X). We ran the following PCR program:

Lid: 105℃, volume: 15 *μ*l

1. 98℃, 30 sec
2. 98℃, 30 sec
3. 62℃, 30 sec
4. 72℃, 30 sec
5. go to step 2 and repeat 11x times
6. 72℃, 5 min
7. 4℃, hold

After validating the correct size of the samples and the absence of amplification in the no-template control, we pooled the libraries together and purified using the Monarch PCR & DNA Cleanup kit (#T1030L), and sequenced on a NovaSeq lane.

#### Barcode extraction pipeline

To extract and count DNA barcodes from Illumina sequencing reads, we first merge the forward and reverse reads using PEAR (Zhang et al. 2014). We then demultiplex the reads further by the indices contained in primers from the first step of PCR. To do this, we used a custom Python script (“process_merged_reads.py”). First, we quality-filtered the reads, discarding any assembled reads with mean Phred quality score across the 20 nucleotide barcode window less than 30. This window corresponds to positions 82-101 of the merged read. The two indices which indicate the sample identity were then extracted by anchoring on the flanking constant regions of the primer (constant motifs ‘GACTATTCTGAT’ and ‘CTTGACGTGC’, each permitting up to 3 mismatches/indels); the downstream index (index 2) was reverse-complemented. Each extracted index was matched to a reference primer table by approximate matching, allowing up to 3 mismatches or indels. We also record a per-index confidence score, which is (1 – (edit distance/index length)). The barcode was captured using a hierarchical cascade of ten fuzzy regular expressions that anchor on the barcode’s constant flanking regions, allowing 2 errors. We capture the internal variable region. We first target a 20 nucleotide intervening region, and include additional patterns because we know that some barcodes differ in length and construction based on previous studies (Dutta et al. 2026). The output is a per-read table recording the sample, the two raw and matched indices with confidence scores, the combined first-step-primer identity, and the extracted barcode. Read parsing was parallelized across allocated CPU cores using Python multiprocessing, processing reads in chunks of 50,000.

To assign each barcode to a strain, we used a second custom Python script called “optimized_barcode_counting.py.” Each read’s inferred sample index combination was validated against a sample sheet specifying the expected first-step-primer pair for every sample; reads were classified as valid matches, index-combination mismatches, sample-number mismatches, or invalid patterns, and these tallies were written to a per-sample diagnostic file. Only reads with a validated, expected index combination were retained. Retained barcodes were matched to a reference barcode-to-strain map (’barcode_to_strain_map_july2026.csv’) supplemented with 6 defined spike-in control barcodes (“SKY” barcodes). These barcoded strains serve as reference strains for the fitness inference. Matching was performed by approximate string matching (RapidFuzz, “ratio” scorer), accepting the best-scoring reference barcode with a similarity score ≥ 90 (out of 100) and leaving reads below this threshold unassigned. Reads were then counted per unique combination of sample, first-step-primer pair, and assigned strain barcode, and the resulting counts were joined back to the sample metadata (experiment, timepoint, environment, and replicate) to produce a per-sample counts table. Per-sample count tables were concatenated into a single run-level table (“all_bc_counts_<RUN>_july2026.csv”) for downstream analysis (“concat_csvs.py”).

The pipeline was implemented in Snakemake (v6.15.5) and executed on a SLURM cluster. Analyses used Python 3 with pandas (v1.1.5), NumPy (v1.19.2), the “regex” module (v2021.9.30), and RapidFuzz (v1.7.1). All reference files (primer table, per-run sample sheets, and barcode-to-strain map) and pipeline code are available at https://github.com/omghosh/longterm-natties.

#### Fitness estimates

We estimate initial fitness according to the maximum-likelihood method described in Ghosh et al. 2026, adapted from Ascensao et al. 2023. This method is described in detail in (Ghosh et al. 2026), and source code is available at https://github.com/omghosh/limiting-functions.

Briefly, we treat the read count of a particular barcode as a negative binomial random variable, and use maximum likelihood inference to estimate the initial frequency of the barcode, and the selection coefficient, or fitness value, from the entire frequency trajectory, under a standard model of frequency change and exponential growth. We infer the noise and standard error on our estimation, as well as the mean fitness of the population, from the reference strain with multiple barcodes. We assume that the log fold change in frequency of a barcode compared to the reference strain, corrected for the changing mean fitness of the population, is equivalent to its relative fitness.

For later time interval fitness estimates, we simply use the log fold change in frequency of individual barcodes between the two time points, which reflects the fitness of these strains relative to the mean of the population.

### Null models of the pooled competition

#### Simulation design

We simulated strain trajectories in each environment 1,000 times. Within each simulation run, we modelled serial batch transfer for 200 cycles. We drew fitness value for each strain from a normal distribution centred on the estimated value with the standard error as its spread, truncated below at s = −4. In null model 1, we drew each fitness value once and held it fixed for the whole simulation run. In null model 2, we redrew fitness independently every transfer. Within a cycle, abundances grew as n_i_ = fᵢ·exp(sᵢ). The transfer bottlenecks were modelled as Poisson sampling: we drew the number of cells transferred from a normal distribution with mean 10^9^ and standard deviation 10^8^, divided by the dilution factor of 250, and simulated post-transfer counts as nᵢ_+1_ ∼ Poisson(N·nᵢ/Σn). Random seeds were fixed for reproducibility. We also reran both null models twice more, using frequency or fitness as the only input.

### Comparison of the experiment with null model

We compared experimental pool composition at transfers 5, 9, 23 and the last sequenced transfer of each environment with the null models. Where the last sequenced transfer differed between replicates, all replicates were assigned to the earlier transfer, but the interpretation does not change we pick a later transfer instead. We calculated the mean frequency of each strain across replicates. We subsampled simulated replicates without replacement to match the number of experimental replicates available for each environment⋅timepoint combination. We repeated the comparison 1000 times with different bootstrapped simulated replicates. To quantify the difference between the experiment and the null models, we used Bray–Curtis dissimilarity: Σ_strains_|f_exp_ − f_sim_| / Σ_strains_(f_exp_ + f_sim_).

#### ‘Finalist’ clone isolation

To isolate the ‘finalists’ of the long-term competition, we first revived the glycerol stocks from the last available timepoint for each media type in 1 mL of YPD media and grew them overnight. Next, we plated the overnight cultures on solid YPD media. When the colonies appeared, we isolated 2 individual colonies from each timepoint of each media type, and used them to grow new overnight cultures. We saved these cultures as glycerol stocks and used them for further DNA extractions and follow-up experiments.

#### ‘Finalist’ clone whole genome sequencing

To sequence the ‘finalists’ of the long-term competition, we first inoculated their glycerol stocks into 1 mL of YPD media and grew overnight. Next day, we spun down the cultures and proceeded with DNA extraction as described above. We then used the transposome tagmentation method (Illumina DNA Prep kit, cat #20060060) to fragment the DNA and amplify libraries for sequencing.

### Whole genome sequencing analysis of the finalists

#### Strain identification

When we isolated finalists from the last timepoints, we initially did not know their strain background. To ID the finalists, we used both barcode and whole genome sequencing information. First, we ‘fished out’ barcode sequences from the WGS reads. We chose the region around the barcode such that it spans outside the region we use for the amplicon sequencing, in order to avoid cross contamination with the amplicon sequencing performed on the same lane.

Next, we designed a custom script to compare genomes from our finalist strains to the genomes of all natural strains sequenced (Loegler et al. 2024). For each of our finalists, we calculated how many of its variants matched with the variants from each of the 3034 sequenced natural strains. We assigned each strain with high (>99% variants matched), medium (> 95% variants matched), or low (> 90% variants matched) match level.

The finalist strains in the SC-HU media, CCD, CMP, and CMQ had very similar genomes. Thus, to differentiate between them, we called variants on all finalists from the SC-HU media. Then, we isolated unique variants that separate one strain from these three strains using the published data. Then, we compared genomes from each of the finalists to the collection of unique variants that can be used to identify the strains between these three.

After validating each strain based on its ‘fished’ barcodes and variant match to the published genomes, we generated .txt files with strains from each of the genetic backgrounds (including ancestors and the evolved clones). We used bwa (H. Li and Durbin 2009) to align the reads to the S288C reference genome and generated gvcf files. We merged all gvcf files into vcf files for all samples with the same ancestral strain based on the generated .txt files. Lastly, we used bcftools to generate a csv file from each vcf file, which allowed us to do downstream filtering and analysis on the called variants. Source code is available at https://github.com/omghosh/longterm-natties.

#### Identifying *de novo* variants

To filter the variants, we first removed all multiallelic variants, the variants that cannot be confidently called in the ancestral strains, the variants with coverage less than 5 or more than double of the median coverage of the sample, and genotype quality of less than 99. For each strain, we called a variant a new heterozygous mutation if its allelic frequency in the ancestor and other strains was < 0.2 or > 0.8, and the call in the ancestor was confidently homozygous. Likewise, we called a variant an LOH variant if its allelic frequency is > 0.2 and < 0.8 in all strains but the strain of interest. To annotate each of the identified variants, we ran SnpEff (Cingolani et al. 2012) command using the R64 version of the *Saccharomyces cerevisiae* S288C genome: *snpEff -dataDir /snpEff_data -v -canon R64-5-1 variants_filtered.vcf* > *variants_filtered.ann.vcf*. We defined LOH tracts as uninterrupted stretches of LOH variants.

### Identifying chromosomal changes

To detect aneuploidies and other large chromosomal abnormalities, we first aligned all of the sequenced reads to the reference genome. Next, we ran the Samtools (Danecek et al. 2021) command ‘*samtools depth -a input_merged.sorted.rg.coord.bam > output_.txt*’ to obtain coverage for each of the bases in the reference genome. For samples with the median coverage of more than six, we calculated median coverage per chromosome, normalized it to the median coverage of the sample, and rounded it to the nearest 0.5. To visualize the genome-wide coverage, we calculated the mean coverage depth for each kilobase of the genome sequence and plotted it against the genome coordinates.

#### Identifying LOH events

For each of the heterozygous ancestor strain (AMA, BEP, BFB, BIE), we assembled the set of heterozygous sites that were present in the published ancestral genome (Peter et al. 2018) and present in our own sequencing of the ancestor strains. Because AMA is triploid, we re-inferred its genotypes from the variant allele frequency (VAF) computed from allelic depths (VAF < 0.1 -> 0/0/0; 0.1 ≤ VAF < 0.5 -> 0/0/1; 0.5 ≤ VAF < 0.9 -> 0/1/1; VAF ≥ 0.9 -> 1/1/1), retaining 0/0/1 and 0/1/1 sites as heterozygous. Variants were annotated with SnpEff as described above. We restricted our analysis to variants of HIGH predicted impact, according to SnpEff annotations, lying in genes that carry exactly one such variant, so that convergence at the gene level is equivalent to convergence at the variant level. BIE was excluded from convergence analysis, as SC-SDS yielded only a single sequenced background.

#### Experiment downsampling

Because we sequenced more than one evolved strain from some replicates, we generated 1,000 bootstrap samples in which one evolved clone was drawn at random from each ancestor⋅medium⋅replicate combination. The same 1,000 samples were used for the experimental data and for every null model.

### Defining null models for LOH convergence

Within each bootstrapped sample, LOH was simulated independently for each evolved strain and matched to that clone’s observed LOH load according to the null model:

● Variant-matched null. Total numbers of simulated LOH variants matched the experiment, but genomic locations were drawn randomly.
● Base pair-matched null. The total number of simulated LOH tracts and the length of each tract in basepairs matched the experiment, but genomic locations were drawn randomly.
● Tract-matched null. The total number of simulated LOH tracts and the length of each tract in variants matched the experiment, but genomic locations were drawn randomly.

After we assigned which variants will undergo an LOH event in our simulation, the direction of LOH was chosen at random. For the diploid ancestors, we assigned conversion to the reference or alternative allele equal probability. For the triploid AMA, the probability reflected median experimental values and was 0.015 for converting to the minor allele and 0.985 to the major allele.

#### Estimating convergence

For each high-impact variant, we counted the number of independent replicates converting to the same allele in the experimental and simulated data. If this number was more than 1, we called this variant convergent. We discarded variants that converted to both states within the same ancestor and media. We tallied convergent genes at three levels: within a single ancestor and media, within an ancestor across the two media, and within a media across ancestors, and repeated this process 1000 times. We then compared experimental and null distributions with a one-sided Mann–Whitney U test, testing the hypothesis that the central tendency of the experiment was higher than of a corresponding null.

#### Cloning of the pRCC-N9 plasmid with the GFP and NatR gene

To create a pRCC-N9 plasmid, we amplified a 4704 bp fragment from the plasmid pRCC-N (Generoso et al. 2016) using primers SK18 and SK67 and a 1869 bp fragment from the plasmid pGS62 (Kao and Sherlock 2008) using primers SK72 and SK73. We purified both fragments using a column purification kit (NEB, cat #T1130L) and digested with DpnI (NEB, cat #R0176S) to eliminate the unamplified plasmid. We then performed a Gibson assembly reaction with both PCR fragments (NEB Cat #E2621S), transformed into *E. coli* (NEB Cat #C3019H) and validated successful clones by sequencing. The plasmid map is available in the supplement.

#### Cloning of a GFP-labeled yeast strain for FACS-based fitness measurement

We GFP-tagged an all-purpose commercial diploid wine strain UCD595. We first amplified a fragment from the plasmid pRCC-N9 using primers JV_P2_For and JV_P2_Rev. The resulting fragment contained the NAT resistance marker, GFP under the *ACT1* promoter, alongside regions of homology to the *HO* gene. We then transformed the UCD595 strain with this fragment. To validate the transformation, we amplified DNA from natR clones with the primers JV_P3_For and JV_P3_Rev (Table S1). We screened three separate transformants for GFP expression in a flow-cytometer (Attune NxT cytometer) and proceeded with one transformant in which >99% of detected cells displayed detectable levels of GFP. This transformant is labelled as JV104.

#### Cytometry-based fitness measurement of the evolved clones

We inoculated the selected wild isolates, their evolved derivatives, and a GFP-labeled strain JV104 from glycerol stocks and grown without shaking in 1 mL of a 96-well plate filled with the measurement media for two 48-h transfers of 10 *μ*l. One day later, we mixed the cells in 1:9 ratio (strain of interest:JV104) by volume and took a 20 *μ*l aliquot for cytometry analysis. A pure JV104 strain and pure version of each of the selected wild isolates was used as a control, and each measurement was performed in two biological replicates. On the second day, we performed a 10 *μ*l transfer into a fresh media. After that we continued sampling 20 *μ*l aliquots for FACS measurement on every odd day and transferring the cells in a fresh media on every even day for 1 to 3 transfers. The collected cells for cytometry analysis were stored in 200 *μ*l of PBS solution (Invitrogen, #AM9624) at 4℃ for up to 5 days. Each sample was analysed for at least 10000 events on a Attune NxT cytometer using the ‘BL1-H’ filter. Singlets were selected by sorting out any particles with a ratio of forward scatter area and height less than 0.9 or more than 1.03. We then computed the percentage of fluorescent cells for our control samples with 100% GFP and 100% dark cells of each of our wild isolated. Since not all of GFP cells fluoresce and some of non-GFP cells exhibit weak fluorescence, we used these control cultures to calibrate the thresholds of the ‘SSC-A’ and ‘BL1-H’ parameters. We then applied these thresholds to each of our samples and calculated the percentage of GFP cells in each of them. By subtracting the percentage of GFP cells from 100 we obtained the fraction of each of our strains at each timepoint in the competition with the JV104 strain. We visualized the results by plotting the trajectories of each strain over time and calculated their relative fitness values by computing the areas under the curves for each strain.

## References

Abreu, Clare I., Shaili Mathur, and Dmitri A. Petrov. 2024. “Environmental Memory Alters the Fitness Effects of Adaptive Mutations in Fluctuating Environments.” Nature Ecology & Evolution 8 (9): 1760–75. 10.1038/s41559-024-02475-9.

Ascensao, Joao A., Keon D. Abedi, Aditya N. Prasad, and Oskar Hallatschek. 2025. “Frequency-Dependent Fitness Effects Are Ubiquitous.” *bioRxiv: The Preprint Server for Biology*, August 21, 2025.08.18.670924. 10.1101/2025.08.18.670924.

Ascensao, Joao A., Kelly M. Wetmore, Benjamin H. Good, Adam P. Arkin, and Oskar Hallatschek. 2023. “Quantifying the Local Adaptive Landscape of a Nascent Bacterial Community.” Nature Communications 14 (1): 248. 10.1038/s41467-022-35677-5.

Blundell, Jamie R., Katja Schwartz, Danielle Francois, Daniel S. Fisher, Gavin Sherlock, and Sasha F. Levy. 2019. “The Dynamics of Adaptive Genetic Diversity during the Early Stages of Clonal Evolution.” Nature Ecology & Evolution 3 (2): 293–301. 10.1038/s41559-018-0758-1.

Breslow, David K., Dale M. Cameron, Sean R. Collins, et al. 2008. “A Comprehensive Strategy Enabling High-Resolution Functional Analysis of the Yeast Genome.” Nature Methods 5 (8): 711–18. 10.1038/nmeth.1234.

Cao, Chunlei, Zhengfeng Cao, Peibin Yu, and Yunying Zhao. 2020. “Genome-Wide Identification for Genes Involved in Sodium Dodecyl Sulfate Toxicity in Saccharomyces Cerevisiae.” BMC Microbiology 20 (1): 34. 10.1186/s12866-020-1721-2.

Chan, Angelina, Michelle Hays, and Gavin Sherlock. 2024. “The Viral K1 Killer Yeast System: Toxicity, Immunity, and Resistance.” *Yeast (Chichester*, England) 41 (11–12): 668–80. 10.1002/yea.3987.

Chao, Lin, and Edward C. Cox. 1983. “COMPETITION BETWEEN HIGH AND LOW MUTATING STRAINS OF ESCHERICHIA COLI.” Evolution; International Journal of Organic Evolution 37 (1): 125–34. 10.1111/j.1558-5646.1983.tb05521.x.

Cingolani, Pablo, Adrian Platts, Le Lily Wang, et al. 2012. “A Program for Annotating and Predicting the Effects of Single Nucleotide Polymorphisms, SnpEff: SNPs in the Genome of Drosophila Melanogaster Strain W1118; Iso-2; Iso-3.” Fly 6 (2): 80–92. 10.4161/fly.19695.

Crabtree, Angela M., Emily A. Kizer, Samuel S. Hunter, et al. 2019. “A Rapid Method for Sequencing Double-Stranded RNAs Purified from Yeasts and the Identification of a Potent K1 Killer Toxin Isolated from Saccharomyces Cerevisiae.” Viruses 11 (1): 70. 10.3390/v11010070.

Danecek, Petr, James K. Bonfield, Jennifer Liddle, et al. 2021. “Twelve Years of SAMtools and BCFtools.” GigaScience 10 (2): giab008. 10.1093/gigascience/giab008.

Desai, Michael M., and Daniel S. Fisher. 2007. “Beneficial Mutation Selection Balance and the Effect of Linkage on Positive Selection.” Genetics 176 (3): 1759–98. 10.1534/genetics.106.067678.

Dunham, Maitreya J., Marc R. Gartenberg, and Grant M. Brown. 2015. Methods in Yeast Genetics and Genomics. Cold Spring Harbor laboratory press.

Dutta, Abhishek, Fabien Dutreux, and Joseph Schacherer. 2021. “Loss of Heterozygosity Results in Rapid but Variable Genome Homogenization across Yeast Genetic Backgrounds.” eLife 10 (June): e70339. 10.7554/eLife.70339.

Dutta, Abhishek, Fabien Dutreux, and Joseph Schacherer. 2022. “Loss of Heterozygosity Spectrum Depends on Ploidy Level in Natural Yeast Populations.” Molecular Biology and Evolution 39 (11): msac214. 10.1093/molbev/msac214.

Dutta, Abhishek, Marion Garin, Victor Loegler, et al. 2026. “Population-Scale Chemical Response Revealed by a Barcoded Yeast Collection.” Nature Communications 17 (1): 6928. 10.1038/s41467-026-73532-z.

Earl, David J., and Michael W. Deem. 2004. “Evolvability Is a Selectable Trait.” Proceedings of the National Academy of Sciences of the United States of America 101 (32): 11531–36. 10.1073/pnas.0404656101.

Ferrare, James T., and Benjamin H. Good. 2024. “Evolution of Evolvability in Rapidly Adapting Populations.” Nature Ecology & Evolution 8 (11): 2085–96. 10.1038/s41559-024-02527-0.

Fisher, Daniel S. 2013. “Asexual Evolution Waves: Fluctuations and Universality.” Journal of Statistical Mechanics: Theory and Experiment 2013 (01): P01011. 10.1088/1742-5468/2013/01/P01011.

Generoso, Wesley Cardoso, Manuela Gottardi, Mislav Oreb, and Eckhard Boles. 2016. “Simplified CRISPR-Cas Genome Editing for Saccharomyces Cerevisiae.” Journal of Microbiological Methods 127 (August): 203–5. 10.1016/j.mimet.2016.06.020.

Gentile, Christopher F., Szi-Chieh Yu, Sebastian Akle Serrano, Philip J. Gerrish, and Paul D. Sniegowski. 2011. “Competition between High- and Higher-Mutating Strains of Escherichia Coli.” Biology Letters 7 (3): 422–24. 10.1098/rsbl.2010.1036.

Gerrish, P. J., and R. E. Lenski. 1998. “The Fate of Competing Beneficial Mutations in an Asexual Population.” Genetica 102–103 (1–6): 127–44.

Gerstein, A. C., A. Kuzmin, and S. P. Otto. 2014. “Loss-of-Heterozygosity Facilitates Passage through Haldane’s Sieve for Saccharomyces Cerevisiae Undergoing Adaptation.” Nature Communications 5 (May): 3819. 10.1038/ncomms4819.

Ghosh, Olivia M., Grant Kinsler, Benjamin H. Good, and Dmitri A. Petrov. 2026. “Genotype-Fitness Mapping of Adaptive Mutants Reveals Shifting Low-Dimensional Structure across Divergent Environments.” PLoS Biology 24 (3): e3003618. 10.1371/journal.pbio.3003618.

Gibson, T. C., M. L. Scheppe, and E. C. Cox. 1970. “Fitness of an Escherichia Coli Mutator Gene.” *Science* (New York, N.Y.) 169 (3946): 686–88. 10.1126/science.169.3946.686.

Gou, Liangke, Joshua S. Bloom, and Leonid Kruglyak. 2019. “The Genetic Basis of Mutation Rate Variation in Yeast.” Genetics 211 (2): 731–40. 10.1534/genetics.118.301609.

Jerison, Elizabeth R., Sergey Kryazhimskiy, James Kameron Mitchell, Joshua S. Bloom, Leonid Kruglyak, and Michael M. Desai. 2017. “Genetic Variation in Adaptability and Pleiotropy in Budding Yeast.” eLife 6 (August): e27167. 10.7554/eLife.27167.

Jiang, Pengyao, Anja R. Ollodart, Vidha Sudhesh, Alan J. Herr, Maitreya J. Dunham, and Kelley Harris. 2021. “A Modified Fluctuation Assay Reveals a Natural Mutator Phenotype That Drives Mutation Spectrum Variation within Saccharomyces Cerevisiae.” eLife 10 (September): e68285. 10.7554/eLife.68285.

Kao, Katy C., Katja Schwartz, and Gavin Sherlock. 2010. “A Genome-Wide Analysis Reveals No Nuclear Dobzhansky-Muller Pairs of Determinants of Speciation between S. Cerevisiae and S. Paradoxus, but Suggests More Complex Incompatibilities.” PLoS Genetics 6 (7): e1001038. 10.1371/journal.pgen.1001038.

Kao, Katy C., and Gavin Sherlock. 2008. “Molecular Characterization of Clonal Interference during Adaptive Evolution in Asexual Populations of Saccharomyces Cerevisiae.” Nature Genetics 40 (12): 1499–504. 10.1038/ng.280.

Kinsler, Grant, Yuping Li, Gavin Sherlock, and Dmitri A. Petrov. 2024. “A High-Resolution Two-Step Evolution Experiment in Yeast Reveals a Shift from Pleiotropic to Modular Adaptation.” PLoS Biology 22 (12): e3002848. 10.1371/journal.pbio.3002848.

Kirschner, M., and J. Gerhart. 1998. “Evolvability.” Proceedings of the National Academy of Sciences of the United States of America 95 (15): 8420–27. 10.1073/pnas.95.15.8420.

Kryazhimskiy, Sergey, Daniel P. Rice, Elizabeth R. Jerison, and Michael M. Desai. 2014. “Microbial Evolution. Global Epistasis Makes Adaptation Predictable despite Sequence-Level Stochasticity.” *Science* (New York, N.Y.) 344 (6191): 1519–22. 10.1126/science.1250939.

Lancaster, Samuel M., Celia Payen, Caiti Smukowski Heil, and Maitreya J. Dunham. 2019. “Fitness Benefits of Loss of Heterozygosity in Saccharomyces Hybrids.” Genome Research 29 (10): 1685–92. 10.1101/gr.245605.118.

Levy, Sasha F., Jamie R. Blundell, Sandeep Venkataram, Dmitri A. Petrov, Daniel S. Fisher, and Gavin Sherlock. 2015. “Quantitative Evolutionary Dynamics Using High-Resolution Lineage Tracking.” Nature 519 (7542): 181–86. 10.1038/nature14279.

Li, Heng, and Richard Durbin. 2009. “Fast and Accurate Short Read Alignment with Burrows-Wheeler Transform.” *Bioinformatics* (Oxford, England) 25 (14): 1754–60. 10.1093/bioinformatics/btp324.

Li, Jing, Simon Stenberg, Jia-Xing Yue, et al. 2023. “Genome Instability Footprint under Rapamycin and Hydroxyurea Treatments.” PLoS Genetics 19 (11): e1011012. 10.1371/journal.pgen.1011012.

Loegler, Victor, Anne Friedrich, and Joseph Schacherer. 2024. “Overview of the Saccharomyces Cerevisiae Population Structure through the Lens of 3,034 Genomes.” G3 (Bethesda, Md.) 14 (12): jkae245. 10.1093/g3journal/jkae245.

Marsit, S., M. Hénault, G. Charron, A. Fijarczyk, and C. R. Landry. 2021. “The Neutral Rate of Whole-Genome Duplication Varies among Yeast Species and Their Hybrids.” Nature Communications 12 (1): 3126. 10.1038/s41467-021-23231-8.

Nguyen Ba, Alex N., Ivana Cvijović, José I. Rojas Echenique, et al. 2019. “High-Resolution Lineage Tracking Reveals Travelling Wave of Adaptation in Laboratory Yeast.” Nature 575 (7783): 494–99. 10.1038/s41586-019-1749-3.

Peter, Jackson, Matteo De Chiara, Anne Friedrich, et al. 2018. “Genome Evolution across 1,011 Saccharomyces Cerevisiae Isolates.” Nature 556 (7701): 339–44. 10.1038/s41586-018-0030-5.

Pieczynska, Magdalena D., Dominika Wloch-Salamon, Ryszard Korona, and J. Arjan G. M. de Visser. 2016. “Rapid Multiple-Level Coevolution in Experimental Populations of Yeast Killer and Nonkiller Strains.” Evolution; International Journal of Organic Evolution 70 (6): 1342–53. 10.1111/evo.12945.

Sharp, Nathaniel P., Linnea Sandell, Christopher G. James, and Sarah P. Otto. 2018. “The Genome-Wide Rate and Spectrum of Spontaneous Mutations Differ between Haploid and Diploid Yeast.” Proceedings of the National Academy of Sciences of the United States of America 115 (22): E5046–55. 10.1073/pnas.1801040115.

Shaw, Alisa E., Mattias N. Mihelich, Jackson E. Whitted, et al. 2024. “Revised Mechanism of Hydroxyurea-Induced Cell Cycle Arrest and an Improved Alternative.” Proceedings of the National Academy of Sciences of the United States of America 121 (42): e2404470121. 10.1073/pnas.2404470121.

Smukowski Heil, Caiti. 2023. “Loss of Heterozygosity and Its Importance in Evolution.” Journal of Molecular Evolution 91 (3): 369–77. 10.1007/s00239-022-10088-8.

Smukowski Heil, Caiti S., Christopher G. DeSevo, Dave A. Pai, Cheryl M. Tucker, Margaret L. Hoang, and Maitreya J. Dunham. 2017. “Loss of Heterozygosity Drives Adaptation in Hybrid Yeast.” Molecular Biology and Evolution 34 (7): 1596–612. 10.1093/molbev/msx098.

Stroud, J. T., B. M. Delory, E. M. Barnes, et al. 2024. “Priority Effects Transcend Scales and Disciplines in Biology.” Trends in Ecology & Evolution 39 (7): 677–88. 10.1016/j.tree.2024.02.004.

Sunshine, Anna B., Celia Payen, Giang T. Ong, Ivan Liachko, Kean Ming Tan, and Maitreya J. Dunham. 2015. “The Fitness Consequences of Aneuploidy Are Driven by Condition-Dependent Gene Effects.” PLoS Biology 13 (5): e1002155. 10.1371/journal.pbio.1002155.

Tenreiro, S., P. C. Rosa, C. A. Viegas, and I. Sá-Correia. 2000. “Expression of the AZR1 Gene (ORF YGR224w), Encoding a Plasma Membrane Transporter of the Major Facilitator Superfamily, Is Required for Adaptation to Acetic Acid and Resistance to Azoles in Saccharomyces Cerevisiae.” *Yeast* (Chichester, England) 16 (16): 1469–81. 10.1002/1097-0061(200012)16:16%3C1469::AID-YEA640%3E3.0.CO;2-A.

Tröbner, W., and R. Piechocki. 1981. “Competition Growth between Escherichia Coli mutL and Mut+ in Continuously Growing Cultures.” Zeitschrift Fur Allgemeine Mikrobiologie 21 (4): 347–49.

Tröbner, W., and R. Piechocki. 1984. “Competition between Isogenic mutS and Mut+ Populations of Escherichia Coli K12 in Continuously Growing Cultures.” Molecular & General Genetics: MGG 198 (2): 175–76. 10.1007/BF00328719.

Verduyn, C., E. Postma, W. A. Scheffers, and J. P. Van Dijken. 1992. “Effect of Benzoic Acid on Metabolic Fluxes in Yeasts: A Continuous-Culture Study on the Regulation of Respiration and Alcoholic Fermentation.” *Yeast (Chichester*, England) 8 (7): 501–17. 10.1002/yea.320080703.

Woods, Robert J., Jeffrey E. Barrick, Tim F. Cooper, Utpala Shrestha, Mark R. Kauth, and Richard E. Lenski. 2011. “Second-Order Selection for Evolvability in a Large Escherichia Coli Population.” *Science (New York*, N.Y.) 331 (6023): 1433–36. 10.1126/science.1198914.

Wu, Xiaohua, and Anna Malkova. 2021. “Break-Induced Replication Mechanisms in Yeast and Mammals.” Current Opinion in Genetics & Development 71 (December): 163–70. 10.1016/j.gde.2021.08.002.

Yone, Hideyuki, Hiromitsu Kono, Hayato Hirai, and Kunihiro Ohta. 2022. “Gene Mapping Methodology Powered by Induced Genome Rearrangements.” Scientific Reports 12 (1): 16658. 10.1038/s41598-022-20999-7.

Zhang, Jiajie, Kassian Kobert, Tomáš Flouri, and Alexandros Stamatakis. 2014. “PEAR: A Fast and Accurate Illumina Paired-End reAd mergeR.” *Bioinformatics* (Oxford, England) 30 (5): 614–20. 10.1093/bioinformatics/btt593.

Zhao, Fengguang, Jiaming Yang, Jingwen Li, et al. 2020. “Multiple Cellular Responses Guarantee Yeast Survival in Presence of the Cell Membrane/Wall Interfering Agent Sodium Dodecyl Sulfate.” Biochemical and Biophysical Research Communications 527 (1): 276–82. 10.1016/j.bbrc.2020.03.163.

Zhu, Yuan O., Gavin Sherlock, and Dmitri A. Petrov. 2016. “Whole Genome Analysis of 132 Clinical Saccharomyces Cerevisiae Strains Reveals Extensive Ploidy Variation.” G3 (Bethesda, Md.) 6 (8): 2421–34. 10.1534/g3.116.029397.

