## Supplemental figures for "Adaptive evolution can overwrite initial natural fitness variation only in highest-fitness yeast isolates"

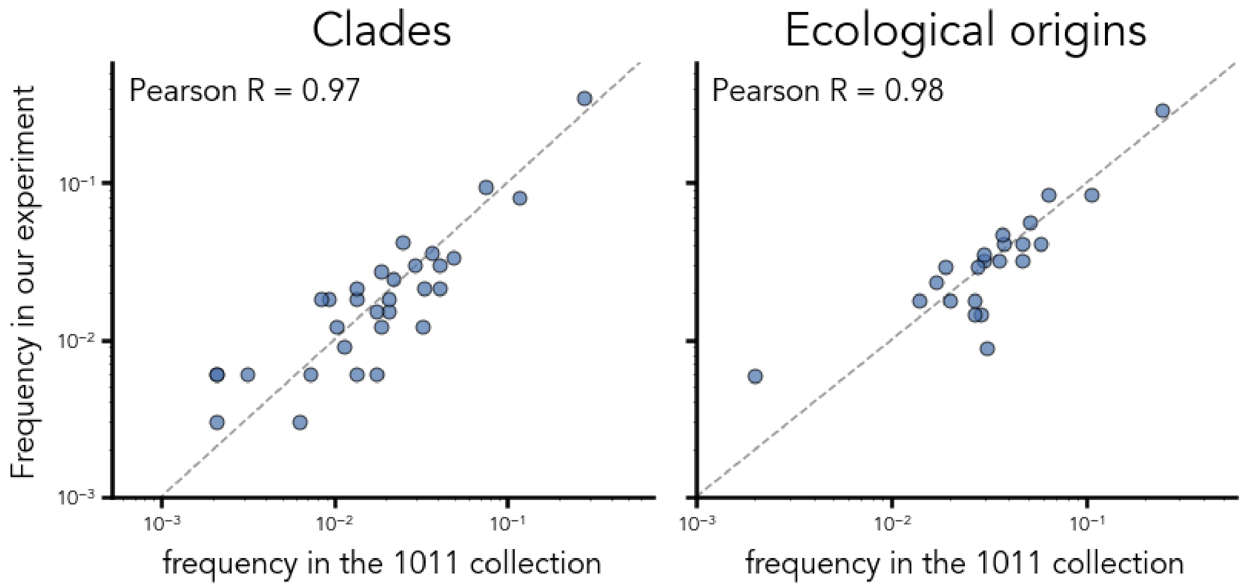


**Fig. S1.** Comparison of the strain collection used in this experiment (282) with the 1011 yeast collection (Peter *et al.,* 2018).

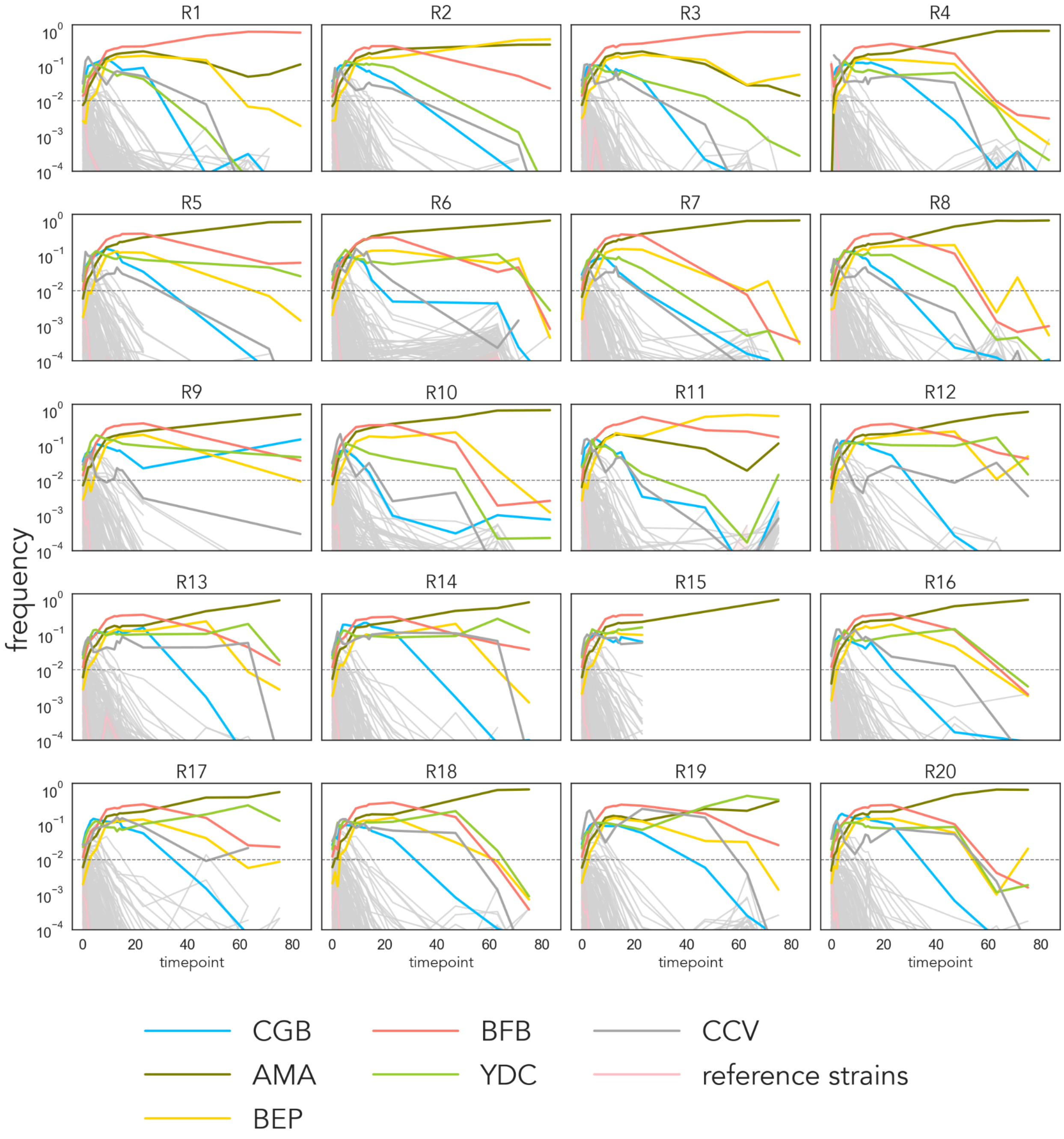


**Fig. S2.** Strain trajectories in the M3 environment.

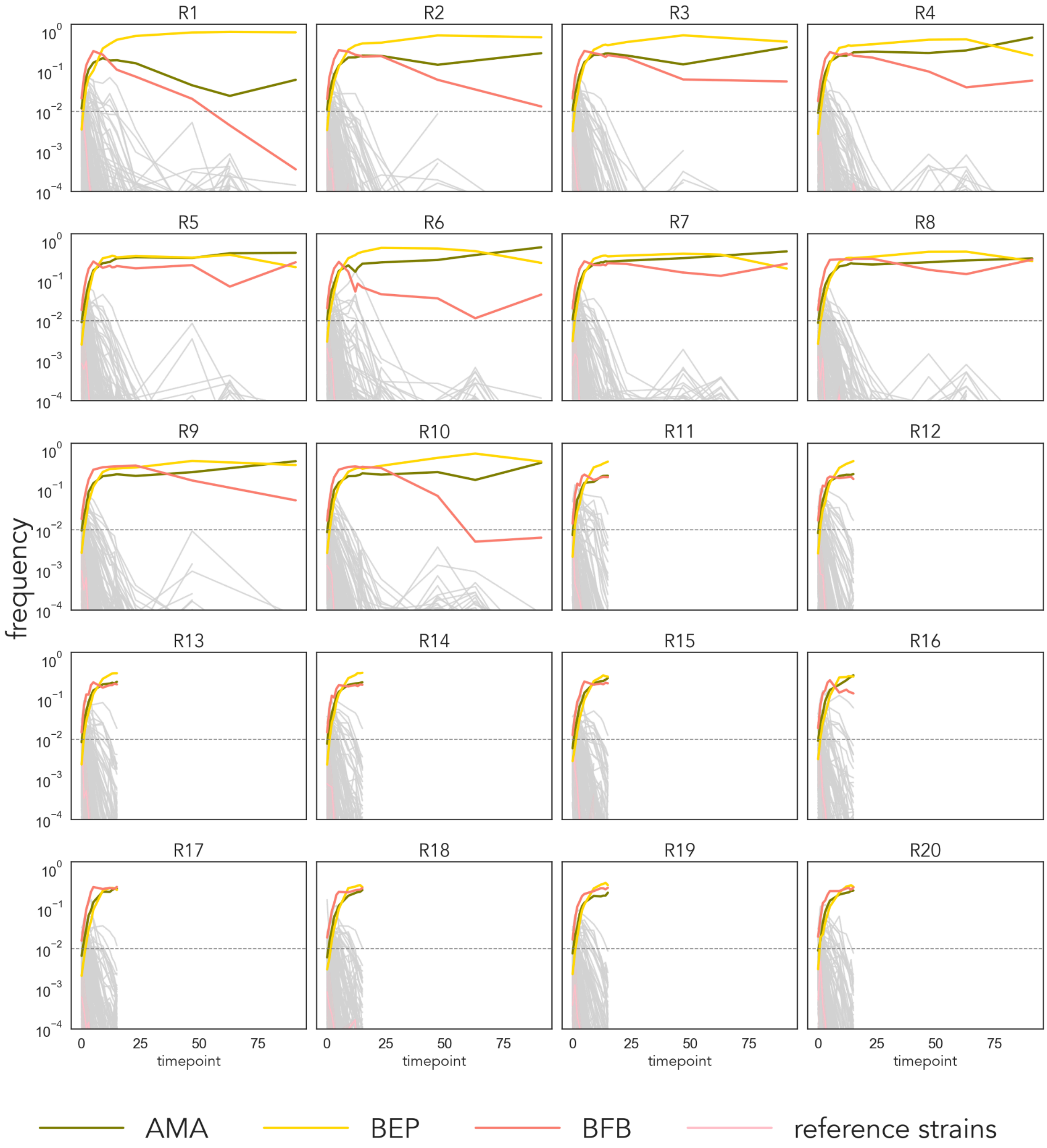


**Fig. S3.** Strain trajectories in the M3-5GLU environment.

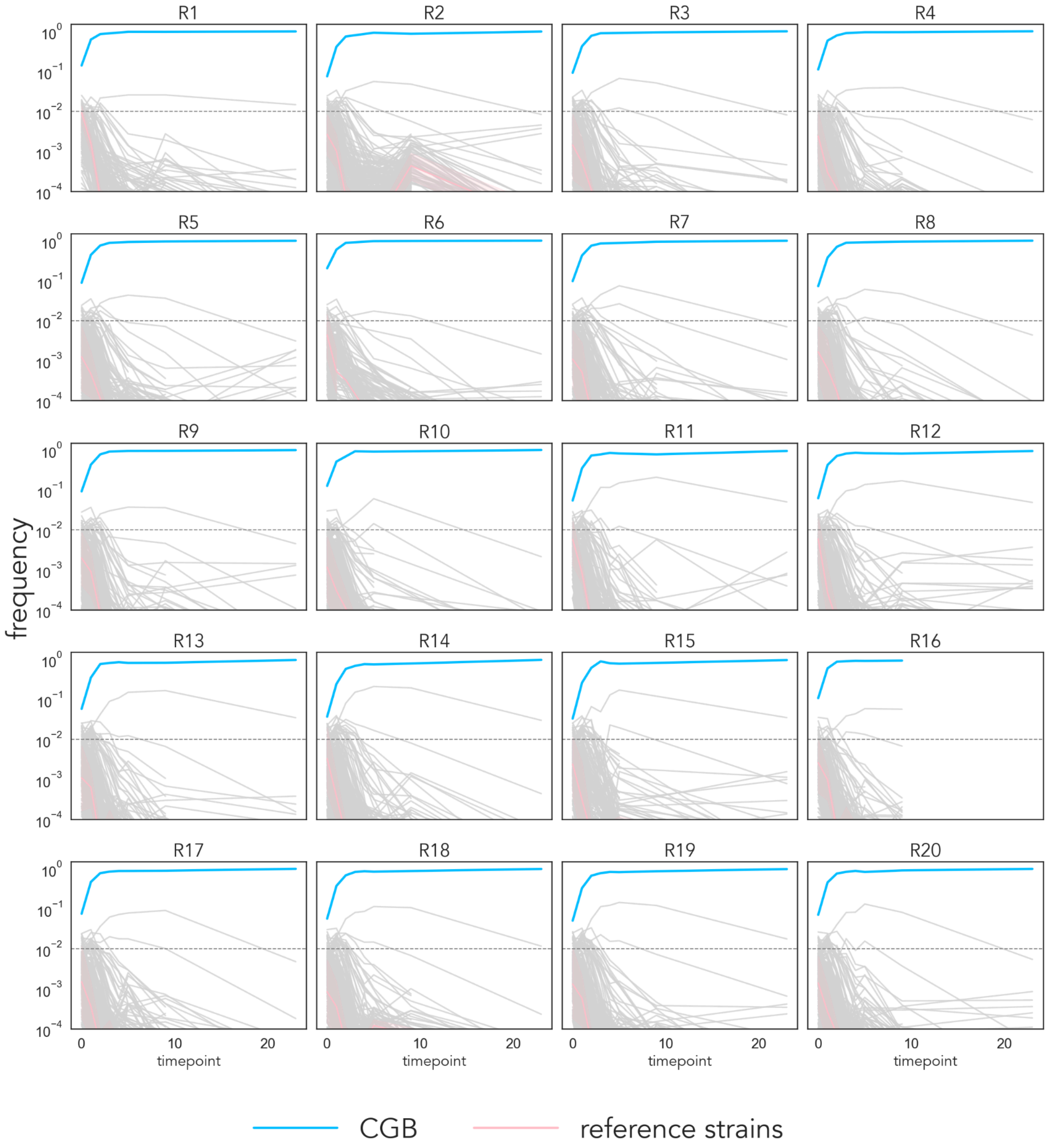


**Fig. S4.** Strain trajectories in the M3-37 environment.

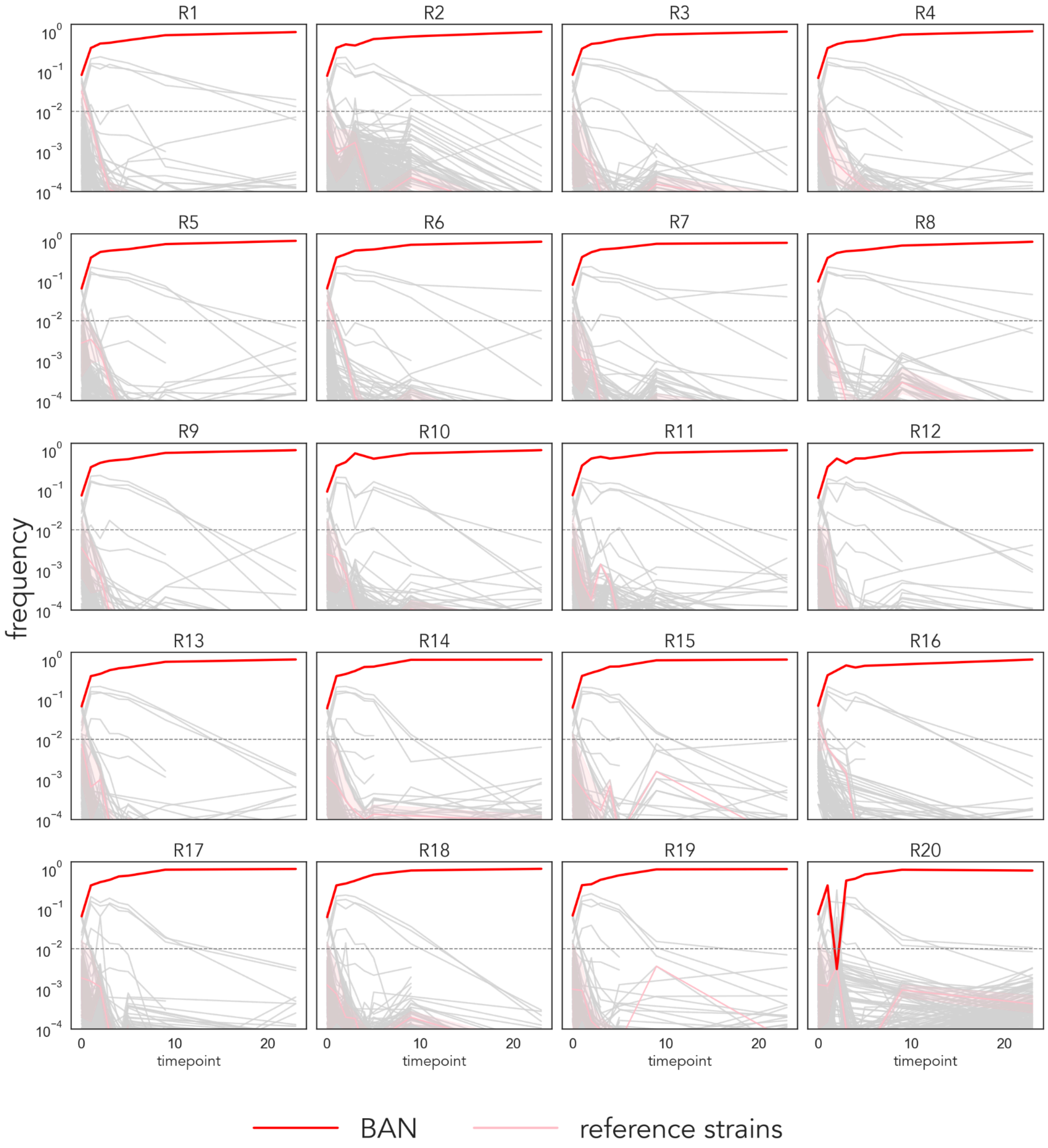


**Fig. S5.** Strain trajectories in the SC environment.

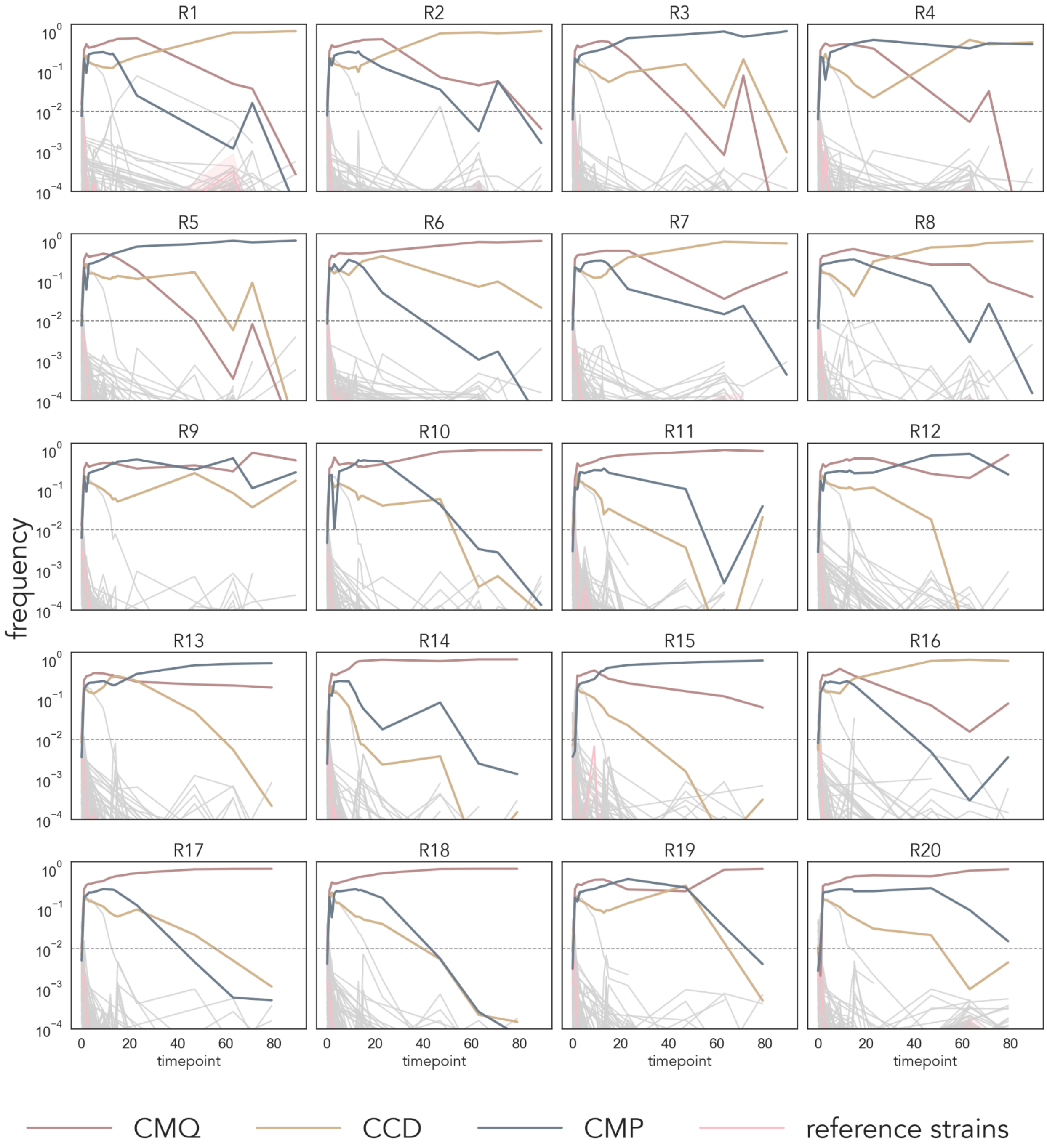


**Fig. S6.** Strain trajectories in the SC-HU environment.

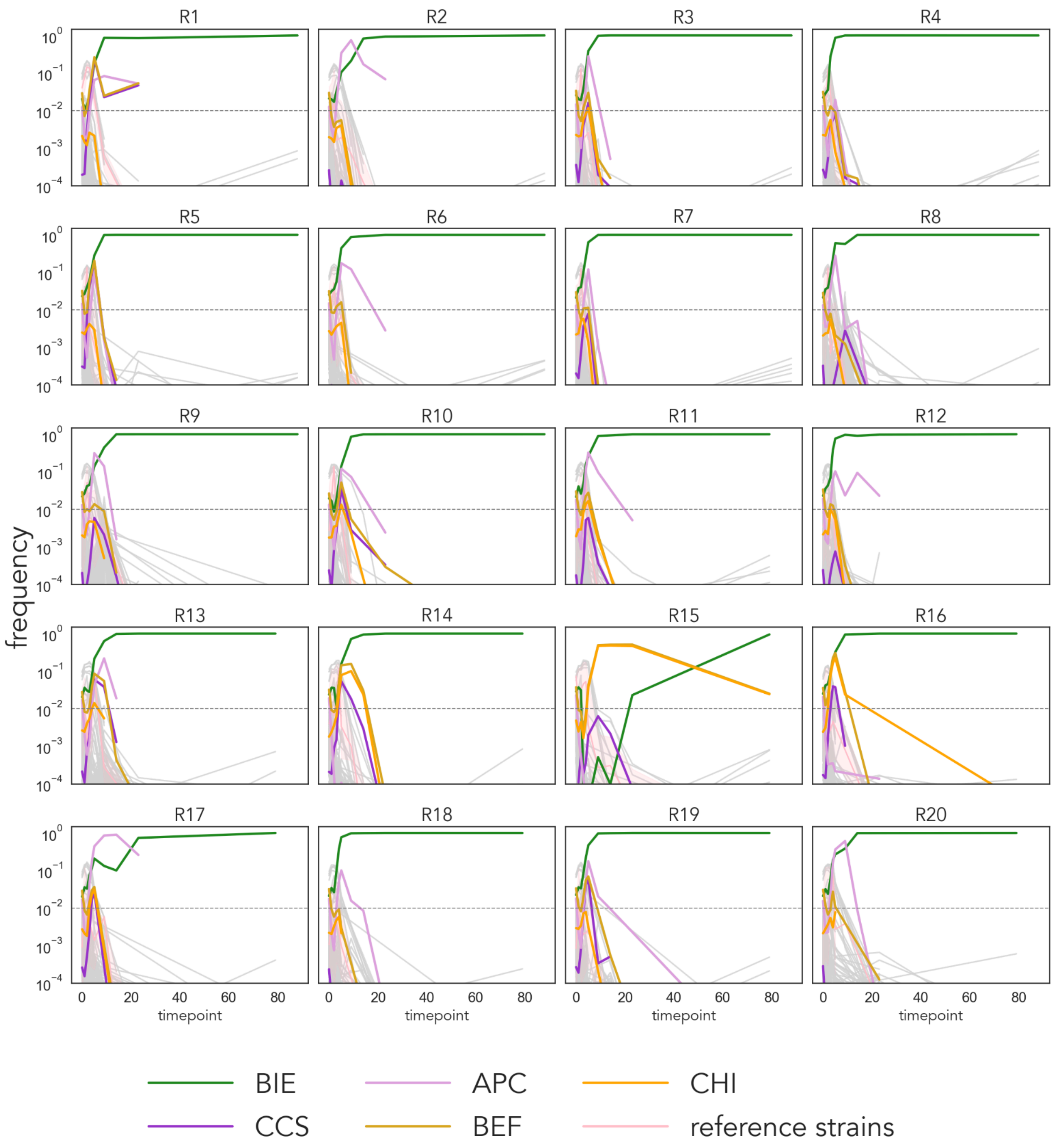


**Fig. S7.** Strain trajectories in the SC-SDS environment. The CCS and APC strains did not persist in any environments, but were included in this figure due to being dominant in the simulated population compositions at the last timepoint.

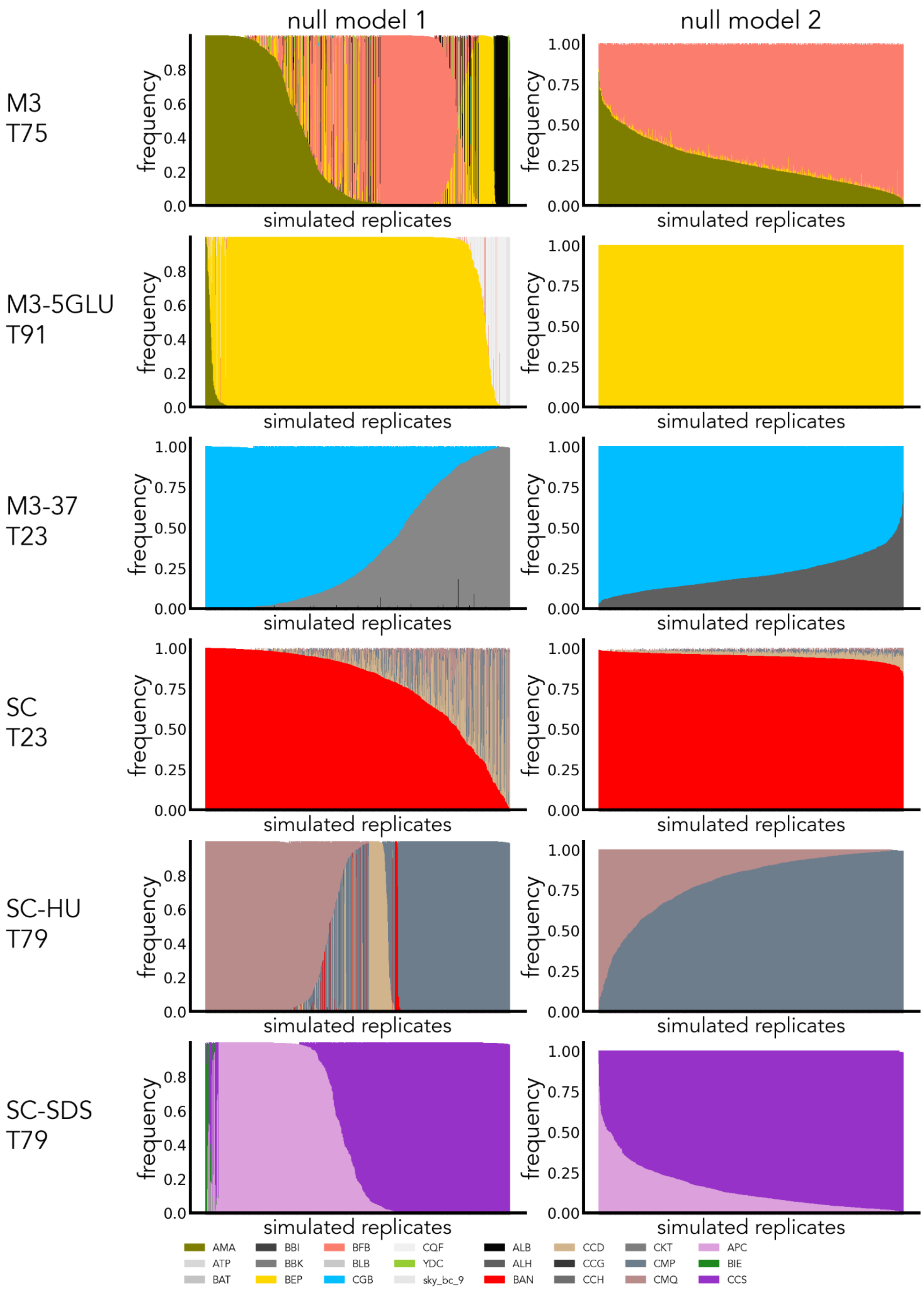


**Fig. S8**. Simulated data for the last experimentally available timepoint for all of our environments and simulation versions 1 (fitness values drawn once per simulation round) and 2 (fitness values redrawn once per cycle).

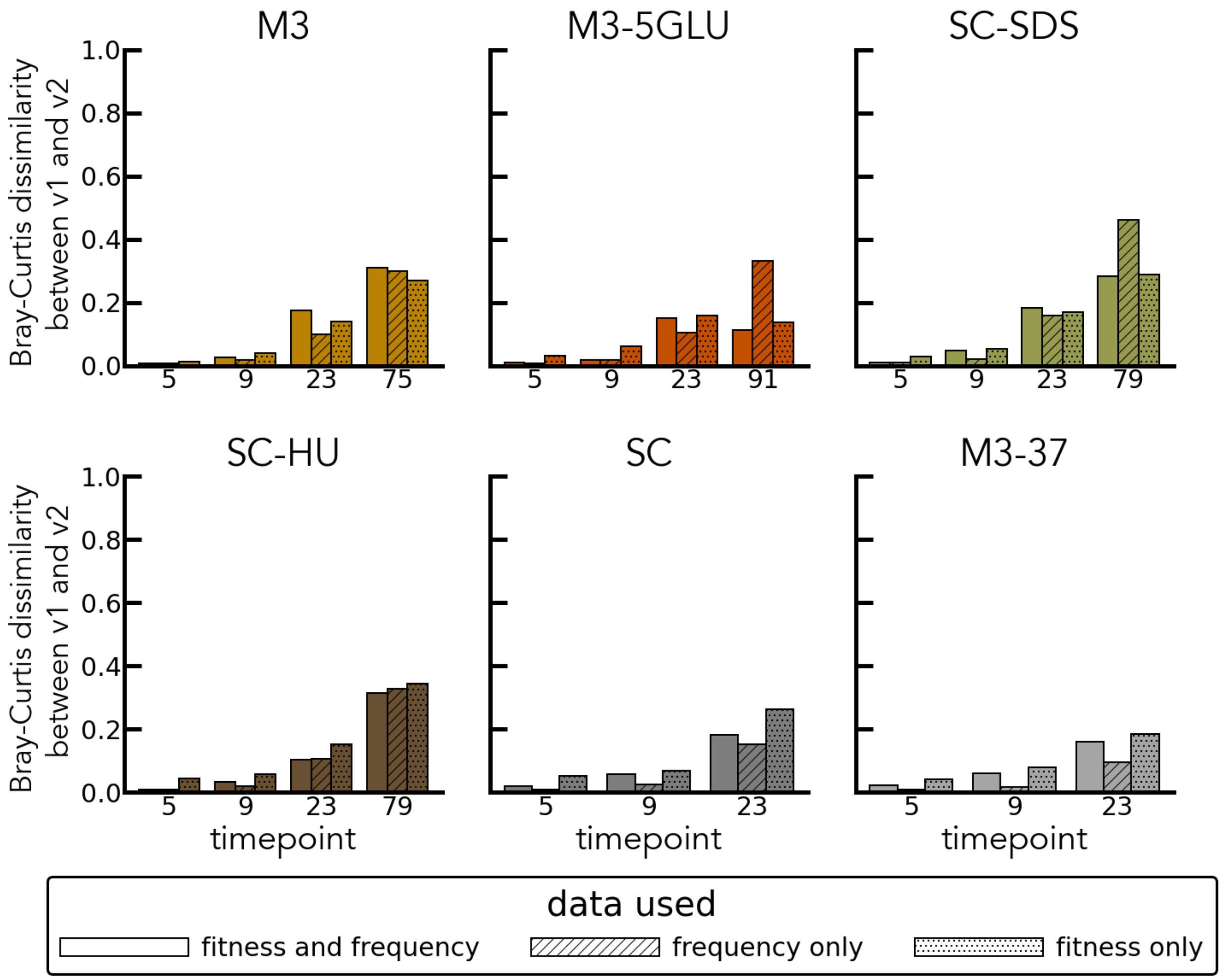


**Fig. S9**. Bray-Curtis dissimilarity between average strains' frequencies between 2 versions of the simulated null models.

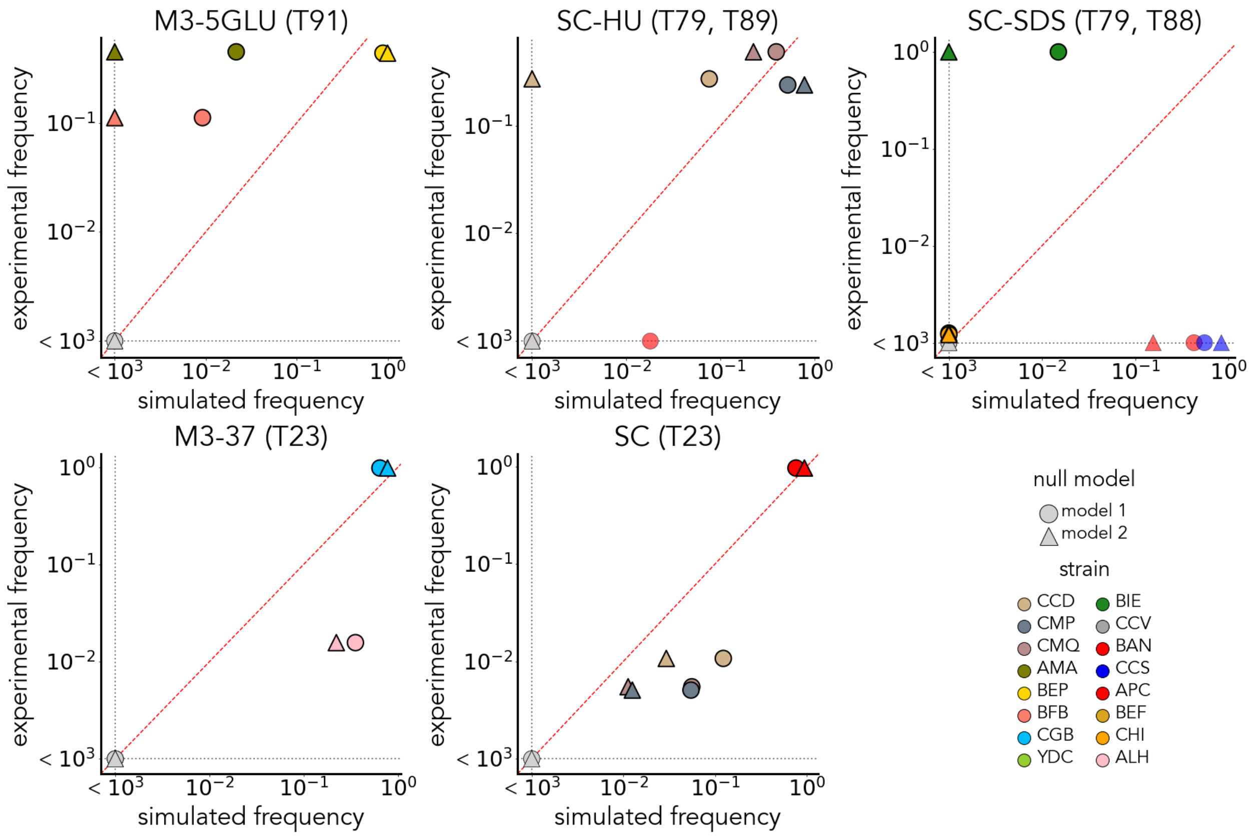


**Fig. S10**. Final frequencies of the top 5% strains in data and simulations (specific timepoints are labeled in the subplots' titles).

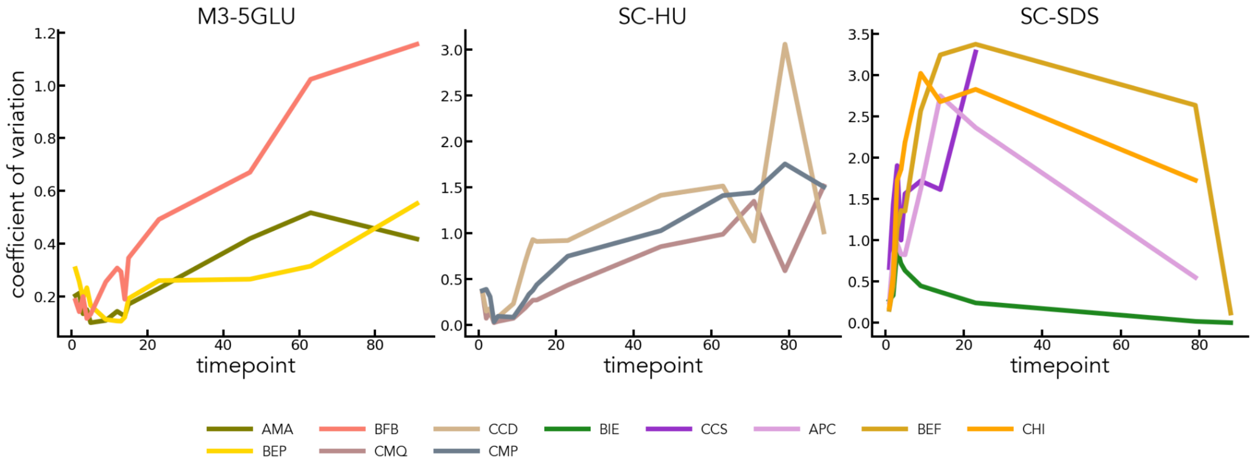


**Fig. S11**. Changes in the coefficients of variation for the finalist strains (as well as CCS and APC in SC-SDS) over time.

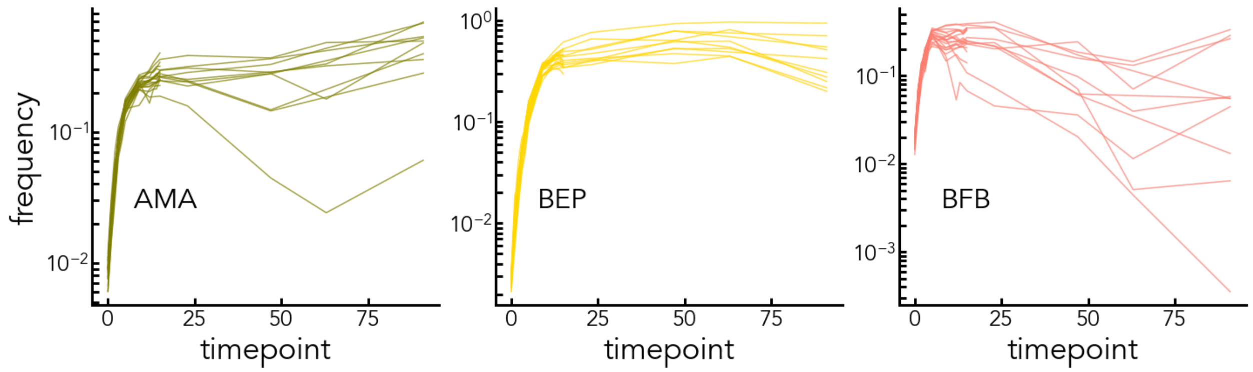


**Fig. S12.** Trajectories of the finalist strains across all experimental replicates in M3-5GLU.

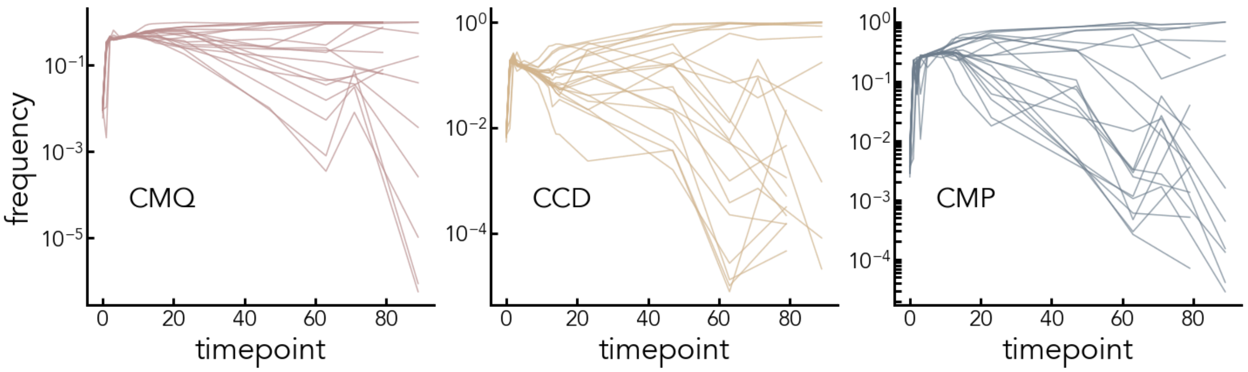


**Fig. S13**. Trajectories of the finalist strains across all experimental replicates in SC-HU.

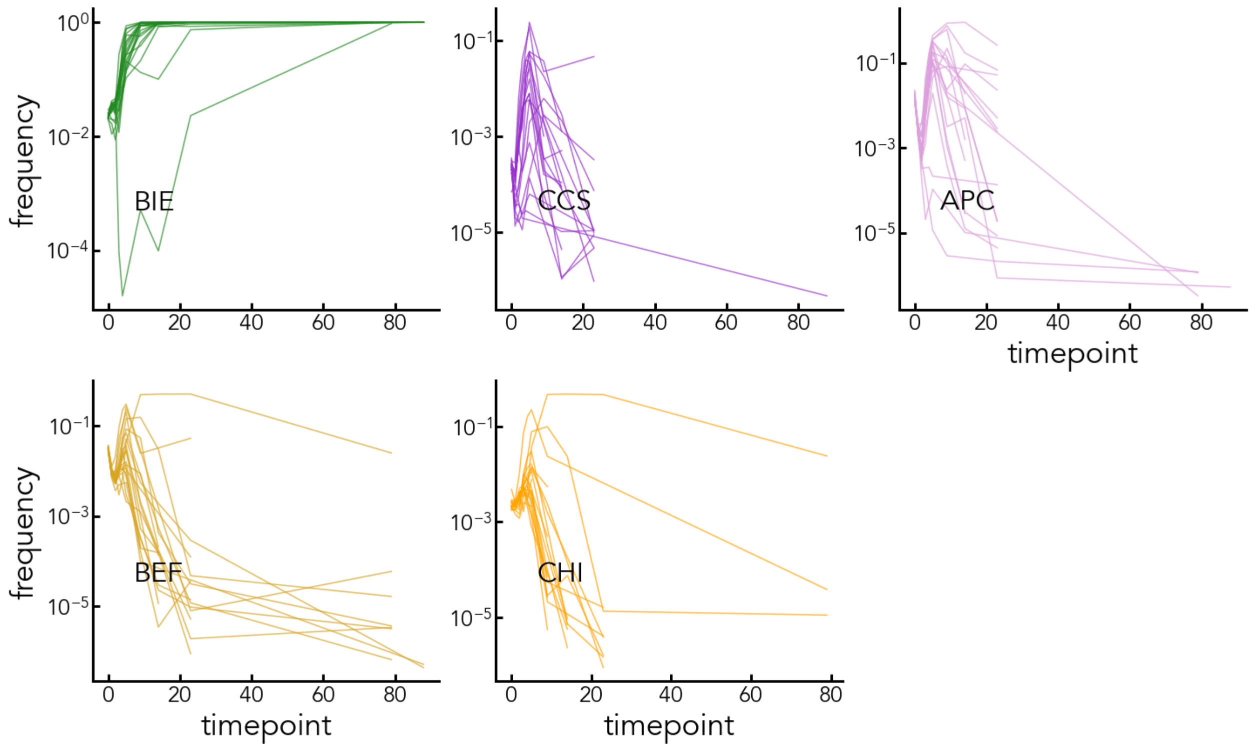


**Fig. S14**. Trajectories of the finalist strains, as well as CCS and APC, across all experimental replicates in SC-SDS.

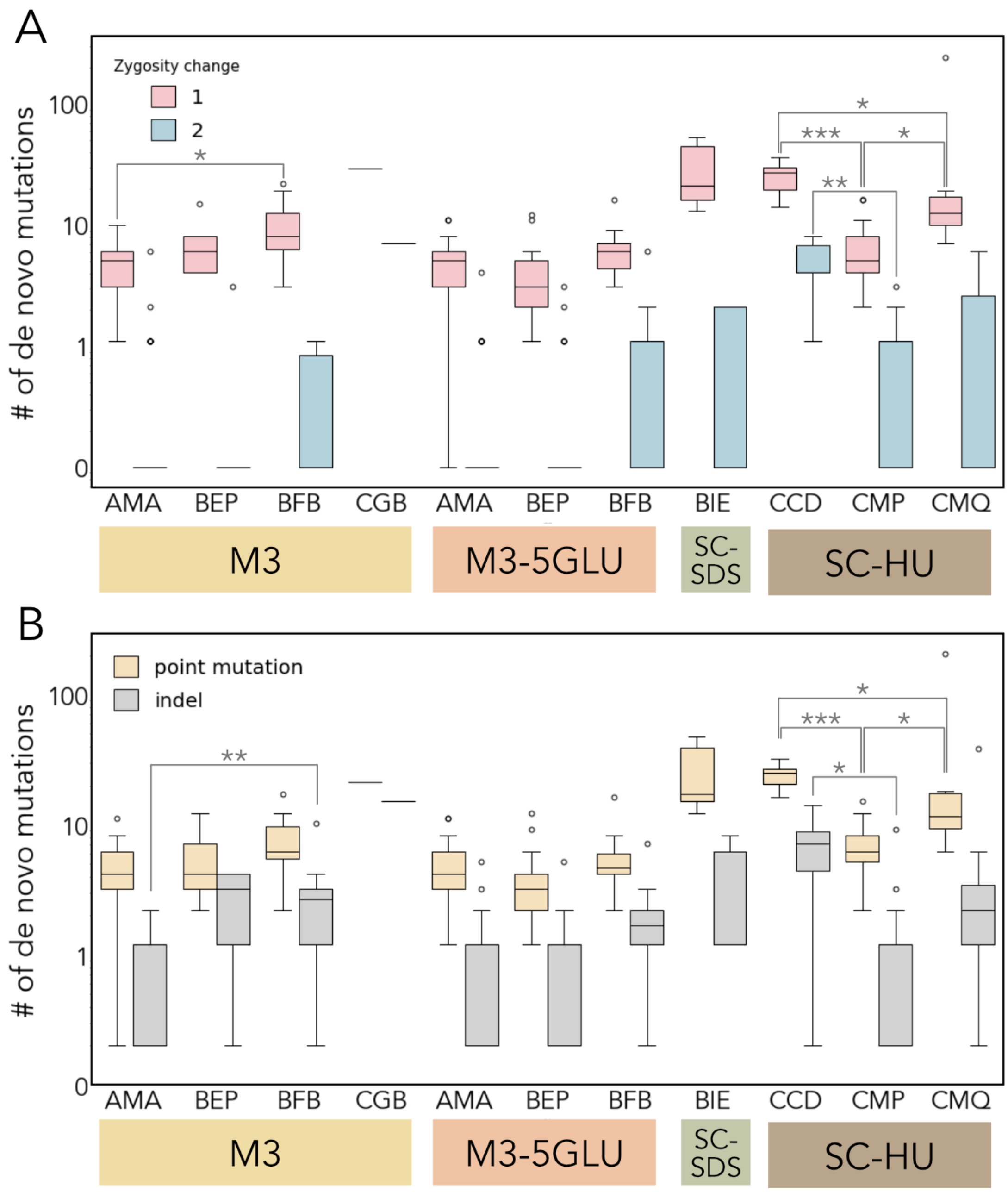


**Fig. S15.** Boxplots showing types of *de novo* mutations by zygosity change (**A**) or molecular nature (**B**) in each media as a function of the strain's genetic background. Asterisks refer to P-values obtained using the Mann-Whitney U test and adjusted for multiple comparisons using the Bonferroni correction. Specifically *: P-value < 0.05, **: P < 0.01, ***: P-value < 0.001. Only statistically significant changes are marked. Only strains derived from the same evolution media were considered for this analysis. In **A**, zygosity change of 1 represents *de novo* heterozygous mutations and zygosity change of 2 represents de novo homozygous mutations for the diploid strain background. For the triploid strain AMA, both zygosity changes of 1 or 2 represent de novo heterozygous mutations. No *de novo* homozygous mutations (zygosity change of 3) were detected in this strain background.

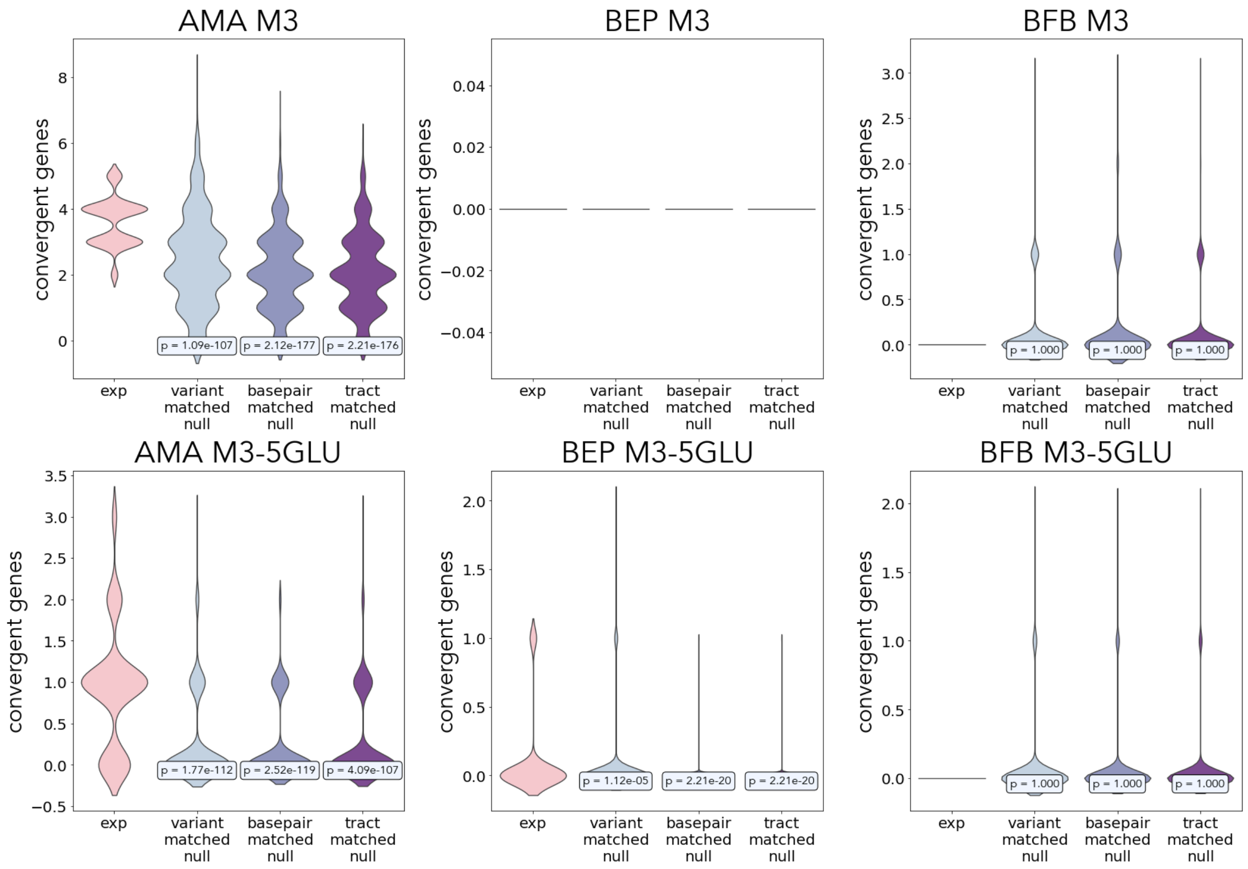


**Fig. S16**. Quantification of convergent genes between replicates of the same media and strain background. For each of 1000 iterations of the calculation, one evolved mutant per replicate per strain background was selected, and the number of genes with an LOH event converting a high-impact heterozygous variant to the same state in more than one independent replicate was calculated. Three null models were used: in Null1, the total number of LOH variants was matched in each strain to the experimentally determined number in this strain; in Null2, the total number of LOH tracts and their length calculated in total base pairs was matched in each strain to the experimentally determined number in this strain; in Null3, the total number of LOH tracts and their length calculated in LOH variants was matched in each strain to the experimentally determined number in this strain. P-values reflect a one-sided Mann-Whitney U test, comparing the experimental distribution with each of the nulls, where an alternative hypothesis is that the central tendency of the null distribution is smaller than in the experimental one.

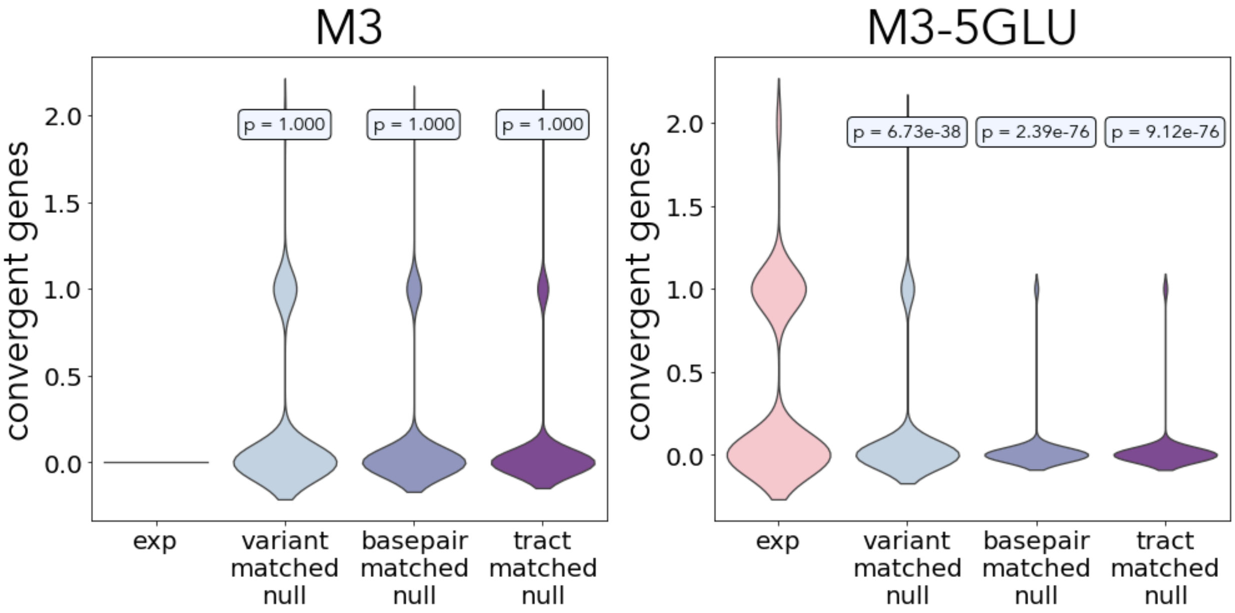


**Fig. S17**. Quantification of convergent genes between replicates from different strain backgrounds evolving in the same media. For each of 1000 iterations of the calculation, one evolved mutant per replicate per strain background was selected, and the number of genes with an LOH event converting a high-impact heterozygous variant to the same state in more than one independent replicate was calculated. Three null models were used: in Null1, the total number of LOH variants was matched in each strain to the experimentally determined number in this strain; in Null2, the total number of LOH tracts and their length calculated in total base pairs was matched in each strain to the experimentally determined number in this strain; in Null3, the total number of LOH tracts and their length calculated in LOH variants was matched in each strain to the experimentally determined number in this strain. P-values reflect a one-sided Mann-Whitney U test, comparing the experimental distribution with each of the nulls, where an alternative hypothesis is that the central tendency of the null distribution is smaller than in the experimental one.

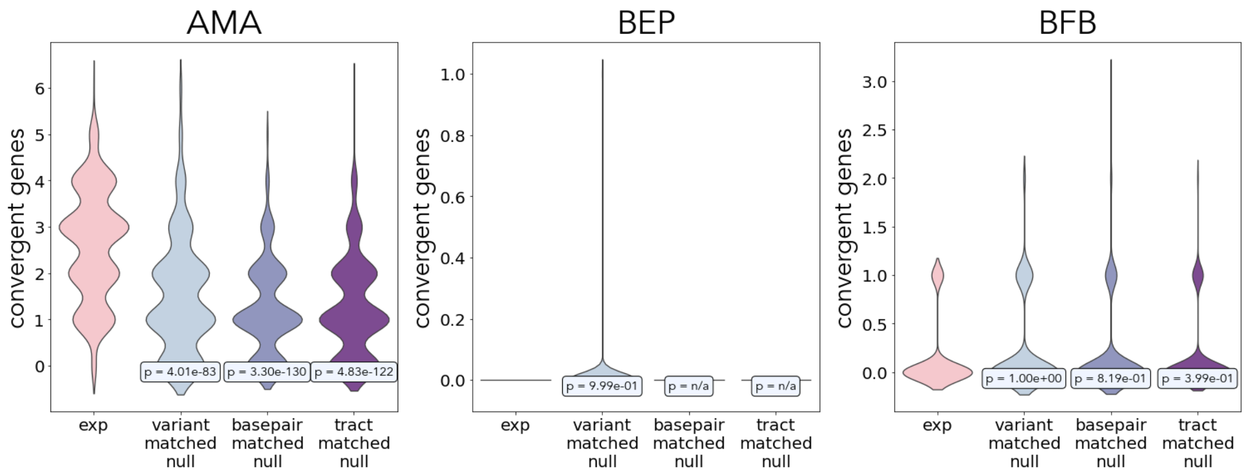


**Fig. S18**. Quantification of convergent genes between replicates of the same strain background evolving in different media. For each of 1000 iterations of the calculation, one evolved mutant per replicate per strain background was selected, and the number of genes with an LOH event converting a high-impact heterozygous variant to the same state in more than one independent replicate was calculated. Three null models were used: in Null1, the total number of LOH variants was matched in each strain to the experimentally determined number in this strain; in Null2, the total number of LOH tracts and their length calculated in total base pairs was matched in each strain to the experimentally determined number in this strain; in Null3, the total number of LOH tracts and their length calculated in LOH variants was matched in each strain to the experimentally determined number in this strain. P-values reflect a one-sided Mann-Whitney U test, comparing the experimental distribution with each of the nulls, where an alternative hypothesis is that the central tendency of the null distribution is smaller than in the experimental one.

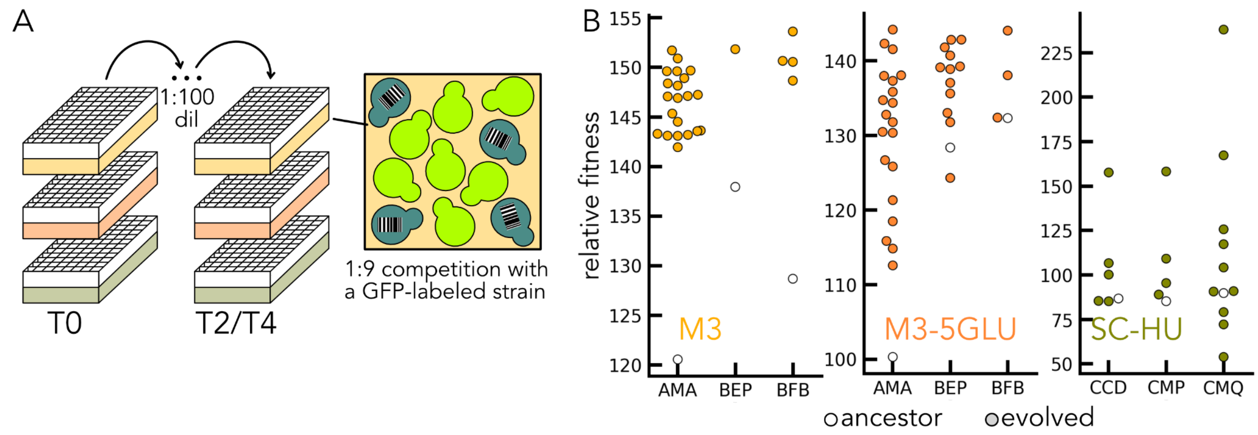


**Fig. S19**. **Functional differences in the evolutionary competition finalists. A.** Individual finalist strains and their corresponding ancestors were cocultured with a GFP-labeled wine strain in 1:9 competitions in 1 mL of media without shaking. We competed the finalists from the M3 and M3-5GLU environments in both of these media, and the finalists from the SC-HU medium in SC-HU. **B.** Relative fitness of each of the strains isolated at the last timepoint relative to its respective ancestor (arbitrary units).
